# Tracing the neural trajectories of evidence accumulation and motor preparation processes during voluntary decisions

**DOI:** 10.1101/2025.10.30.685712

**Authors:** Lauren C. Fong, Paul M. Garrett, Philip L. Smith, Robert Hester, Stefan Bode, Daniel Feuerriegel

## Abstract

Voluntary decisions have previously been described by *where* they arise in the brain and *how* actions corresponding to one’s choice are prepared. However, the processes by which these internally guided decisions are formed and translated into action remain unclear. We tested whether neural signatures of evidence accumulation and motor preparation – established in perceptual decision-making – are also observed during voluntary decisions. Forty-nine adults made voluntary (free choice between two options) and forced (single specified choice) decisions while electroencephalography was recorded. To isolate decision formation from action selection, all responses were made with a right-handed keypress. After applying signal deconvolution and current source density transformations, we examined three signals: the centro-parietal positivity (CPP), Mu/Beta (8 – 30 Hz) amplitudes, and a left-hemisphere readiness potential contralateral to the response hand. For both voluntary and forced decisions, CPP and Mu/Beta amplitudes showed hallmark accumulation-to-bound dynamics – faster responses had steeper ramping signals that converged to stereotyped pre-response amplitude levels. The left-hemisphere readiness potential likewise showed convergence to a pre-response level but with less reliable evidence for RT-dependent slope scaling, resembling a late, effector-specific motor gate rather than an evolving decision variable. Waveform amplitudes across the time course of these signals also did not differ reliably between voluntary and forced decisions. Our results indicate that, like perceptual decisions, voluntary choices are also formed by a graded accumulation process that culminates in a motor action, with the CPP indexing an evolving decision variable and Mu/Beta indexing effector-specific motor readiness.

## 1 Introduction

Every day, we make countless voluntary decisions. Some are trivial – like choosing which mug to drink coffee from – and others more consequential – such as whether to speak publicly about a societal issue or when to tackle a difficult task. Voluntary decisions require us to first recognise that more than one course of action exists – two coffee mugs – then evaluate and deliberate between options by gathering evidence towards each alternative. This process is guided by internally generated goals and preferences (e.g., I prefer the blue mug over the green mug, or I need the insulated mug to keep my coffee warm) and to a lesser extent, by situational context and external cues (Haggard, 2008).

Once sufficient evidence has been accumulated to reach a decision, we commit to a choice by executing the corresponding motor action (Bode et al., 2014; Bogler et al., 2023; Brass et al., 2019; Roskies, 2010; Schurger et al., 2012). In this article, we detail the neural dynamics underlying decisional evidence accumulation in voluntary decision making.

Research on the neurophysiology of volition typically contrasts voluntary (i.e., free-choice) trials with externally cued control trials to isolate the neural processes underlying internally guided decisions. In voluntary trials, one option is selected from among many; in control trials, only one option is available (Brass & Haggard, 2008; Zapparoli et al., 2017). Functional MRI studies consistently show that voluntary trials engage mesial frontal regions – including the pre-supplementary motor area (pre-SMA; e.g., Fried et al., 2011; Passingham et al., 2010; Seghezzi et al., 2019, 2025), supplementary motor area (SMA; e.g., Hoffstaedter et al., 2013; Jahanshahi et al., 1995), and anterior cingulate cortex (e.g., Wiese et al., 2005; Zapparoli et al., 2018) – and parietal areas (e.g., Cunnington et al., 2002; Desmurget et al., 2009; Wisniewski et al., 2016; see Haggard, 2019 for a review of fMRI studies). Multivariate decoding analyses have further demonstrated that choice-selective information can be detected in these areas prior to action, predicting object choices (Bode et al., 2013; Löffler et al., 2020) and arithmetic operations (Haynes et al., 2007; Soon et al., 2013).

Work using electroencephalography (EEG) further provides millisecond-level resolution of motor preparation preceding voluntary actions. One key event-related potential (ERP) component is the readiness potential (RP). This is a slow, negative-going wave measured over central electrodes (e.g., Cz) that gradually increases in amplitude prior to movement (Deecke et al., 1969; Kornhuber & Deecke, 1965) and is thought to originate in the pre-SMA/SMA (Cui et al., 1999; Fried et al., 2011). The RP is larger in amplitude for spontaneous compared to instructed actions (e.g., Dirnberger et al., 2008; Jahanshahi et al., 1995), deliberate versus arbitrary actions (e.g., Masaki et al., 1998; Travers et al., 2021), and when decisions were made with less reliance on exogenous information (e.g., Gluth et al., 2013; Travers & Haggard, 2021). Consequently, the RP has been widely interpreted as an index of motor preparation for voluntary actions.

Despite clear evidence of where voluntary actions are implemented in the brain, the processes of decision formation that precede voluntary action remain underspecified. Classic EEG studies have emphasised action execution markers, like the RP, without characterising the evolving decision dynamics that precede motor implementation, particularly when stimulus-response mappings or effector demands vary across conditions. This is in direct contrast to perceptual forced-choice decision-making studies that have identified neural signals that reflect evidence accumulation dynamics (Kelly et al., 2021; Steinemann et al., 2018), as formalised in sequential sampling models such as the Diffusion Decision Model (DDM; Ratcliff, 1978; Ratcliff and Smith, 2004).

Sequential sampling models jointly explain choice and response times (RTs) using psychologically interpretable parameters: decisions are reached after noisy evidence accumulates over time and crosses a decision-boundary, triggering a motor response (Ratcliff et al., 2016; Smith & Ratcliff, 2004). Hypothesised neural correlates of evidence accumulation trajectories exhibit two defining properties: (i) build-up rates that scale inversely with RTs, often mapping onto the evidence quality (drift rate) used to inform the decision (O’Connell & Kelly, 2021; O’Connell et al., 2012), and (ii) pre-response amplitudes that converge at a common level, consistent with a fixed decision boundary parameter (Kelly & O’Connell, 2013; Philiastides et al., 2014).

The centro-parietal positivity (CPP) has been identified as a neural correlate of an evolving decision variable that exhibits both of these properties. The CPP is a positive-going ERP component measured over midline posterior electrodes (e.g., Pz) that ramps towards stereotyped amplitude levels prior to a response, regardless of RT (Kelly & O’Connell, 2013; O’Connell et al., 2012). This ramping is steeper for faster than for slower decisions (O’Connell & Kelly, 2021; O’Connell et al., 2012), consistent with how higher rates of evidence accumulation produce faster responses. CPP dynamics are dissociable from both sensory encoding and motor preparatory activity, making the CPP a relatively direct, supramodal neural correlate of decision formation (O’Connell & Kelly, 2021; Steinemann et al., 2018). While the CPP has been robustly established in perceptual tasks (e.g., Feuerriegel et al., 2021; Kelly & O’Connell, 2013; Kelly et al., 2021; Steinemann et al., 2018; Twomey et al., 2015), its generalisation beyond perceptual decision-making remains under active debate (Frömer et al., 2024). Despite being previously reported in value-based (Pisauro et al., 2017) and memory-based decisions (van Ede & Nobre, 2024; van Vugt et al., 2019), methodological considerations have prompted caution about interpreting the CPP as a domain-general evidence accumulation decision variable (for further discussion, see O’Connell et al., 2025).

Mu/Beta (MB; 8–30 Hz) spectral amplitudes also reflect accumulation-to-bound dynamics in decision-making tasks where motor responses are required. MB amplitudes gradually decrease prior to an action at electrodes contralateral to the response hand (e.g., C3 and C4), indexing the progressive release of motor inhibition (Donner et al., 2009; Lange et al., 2013; Pfurtscheller & Lopes da Silva, 1999). Like the CPP, the rate of amplitude change scales with RT and reaches a fixed amplitude prior to a motor response (Donner et al., 2009; Lange et al., 2013; O’Connell et al., 2012). Contrary to the CPP, the dynamics of MB activity are sensitive to time constraints and urgency (Kelly et al., 2021; Murphy et al., 2016; Steinemann et al., 2018), consistent with adaptive changes in motor readiness that are responsive to different task demands. Collectively, these findings support MB amplitudes as a motor preparatory signal that accumulates towards a motor-execution boundary and adapts to task dynamics.

Beyond its classic descriptor as a marker of voluntary actions, the RP has been conceptualised within an evidence accumulation framework for simple self-paced actions, modelled as a leaky stochastic accumulator (Schurger, 2018; Schurger et al., 2012). Based on this view, stochastic moment-to-moment fluctuations of neural noise in motor circuits act as “evidence” that ramps toward a motor execution boundary. Once these fluctuations happen to cross the boundary, an action is initiated – analogous to motor threshold-crossing dynamics shown in MB amplitudes described earlier. Under ambiguous or absent external evidence, or in arbitrary decisions with no strong reasons to pick one option over another, threshold-crossing would be primarily noise-driven, and facilitated by lowered boundaries (Brass et al., 2019; Maoz et al., 2019).

This threshold-crossing notion has been extended from arbitrary, spontaneous voluntary actions to more meaningful voluntary decisions. The conditional intention and integration to bound (COINTOB) model proposed by Brass et al. (2019) conceptualises the RP as a neural signature of a general accumulation-to-bound process that is dependent on evidence inputs and policies. In arbitrary, self-paced decision contexts, the accumulator would be dominated by stochastic neural fluctuations; in deliberate decisions, the same evidence accumulation mechanism integrates decision-relevant endogenous (e.g., preferences or goals) and/or external evidence (e.g., monetary values or visual choice stimuli), alongside some noise. The COINTOB model also accommodates lowered or collapsing bounds in decisions made under speed pressure – dynamics similarly observed in MB amplitudes under urgency manipulations. Thus, differences across decision contexts may reflect changes in inputs and bound policies, yet underlie similar evidence accumulation mechanisms.

Empirically, this notion has been supported by recent work showing that single-trial RP durations predict model-derived decision times in voluntary actions (Lui et al., 2021). Racing-accumulator model fits have shown that arbitrary decisions are well captured by noise-driven accumulation, but less so for deliberate decisions, which were better captured by a value-assessment component (Maoz et al., 2019). A lateralised variant of the RP (LRP), derived from the contralateral-ipsilateral difference over motor electrodes (Eimer, 1998; Haggard, 2009), likewise shows accumulation-to-bound dynamics that can scale with sensory evidence in motion discrimination tasks and lags behind the CPP (Kelly & O’Connell, 2013). The motor system is also thought to receive and track evolving decision evidence during deliberation to prime motor preparation and reduce computational costs when an action is required (Selen et al., 2012). Together, these observations make it plausible that RP/LRP can, in some contexts, index decision formation within the motor system, complementing the CPP and MB signals.

Taken together, the CPP, MB, and L/RP have been established as neural correlates of decision evidence accumulation (CPP) and motor execution dynamics (MB, L/RP) in perceptual decision-making. However, unlike in perceptual decisions, voluntary decisions are primarily endogenously driven, lack objectively correct outcomes, and often unfold without strict deadlines or explicit speed-accuracy constraints. Although these distinctions do not necessarily imply differences in neural correlates of evidence accumulation and motor execution between perceptual and voluntary decisions, they raise a critical question: do these signals extend to voluntary decisions and, if so, do they also exhibit canonical accumulation-to-bound properties?

We address these open questions by recording EEG from a large sample of participants (N=49) who completed a colour decision task. We included voluntary decision trials (in which participants chose between two presented options) and forced decision trials (in which participants chose the single presented option). We operationalised voluntary decisions as choices between two concurrently presented alternatives, without an externally specified or objectively correct outcome. As a result, participants not only identified the two-colour options perceptually, but also used endogenous information, such as preferences, goal-related strategies, or other internal heuristics to guide their choices. By contrast, forced trials required a perceptual decision: participants identified and committed to the colour of a single option. Participants indicated the moment they committed to a voluntary or forced colour choice with a right-handed keypress. This design let us focus on the left-hemisphere readiness potential (LHRP) as a contralateral motor-preparatory measure while keeping action selection and stimulus-response mapping the same across choice conditions. It also allowed us to assess voluntary decision dynamics in the absence of concurrent action selection. The right-handed keypress indexed decision commitment (i.e., RT) and was used to response-lock our EEG analyses.

We then tested whether the three candidate signals – CPP, MB, and LHRP – exhibited the two canonical characteristics of an evolving decision variable in voluntary decisions: (a) build-up rates that scale inversely with RTs, and (b) pre-response amplitudes that build to a fixed level, irrespective of RTs (Kelly & O’Connell, 2013). To disentangle EEG signals time-locked to stimulus onset that may overlap with non-stimulus-locked activity (addressing concerns in Frömer et al., 2024), we used Residue Iteration Decomposition (RIDE) to deconvolve different contributions to EEG waveforms (Ouyang et al., 2015). We also applied current source density (CSD) transformations to enhance spatial precision and reduce volume conduction. This allowed us to characterise whether these candidate signals trace the trajectories of an evolving decision variable in more endogenously driven voluntary decisions under tightly controlled motor demands. More specifically, this allowed us to test whether the CPP and MB signals generalise across decision types. We could further compare these dynamics to forced decisions, where this endogenous deliberation was not required. Establishing how these signals behave in voluntary decisions, and how they compare to forced decisions, can provide insights that could inform future joint modelling of EEG measures and behavioural task performance data.

## 2 Method

### 2.1 Participants

We recruited 50 participants (17 men and 33 women) aged between 18-39 years (*M_age_* = 20.88). All participants had normal or corrected-to-normal vision and reported no history of neurological or psychiatric disorders; 42 participants were right-handed. We excluded one participant due to excessively noisy EEG data, leaving 49 participants for analyses. Each participant received either $30 AUD or two course credits. The University of Melbourne Human Research Ethics Committee approved this study (Approval Number 27017).

### 2.2 Stimuli and Apparatus

We presented single- and dual-coloured balloons against a black background (Figure 1). Four colours - blue, green, orange, and pink - were chosen from a CIELAB colour wheel (Wyszecki & Stiles, 1982) sampled 90*^◦^* apart, at 43*^◦^* (orange), 133*^◦^* (green), 223*^◦^* (blue) and 313*^◦^* (pink). They were then converted to RGB values: orange = [206, 104, 0], green = [19, 144, 0], blue = [0, 176, 238], and pink = [236, 0, 216]. The wheel was centred at L = -54, a = 18, b = -8 with a radius of 108.82 mm (as used in Adam et al., 2017), to ensure approximately equal luminance and distinct colours to reduce saliency biases in participants’ choices. Stimuli were presented using Psychtoolbox (Brainard, 1997) interfacing MATLAB R2022b (Mathworks Inc.) on a 24.5-inch ASUS ROG Swift PG258Q monitor (1920 x 1080 pixels, 60 Hz refresh rate). Participants responded using a Keychron Q0 QMK Custom Number Pad (1000 Hz polling rate).

**Figure 1.**
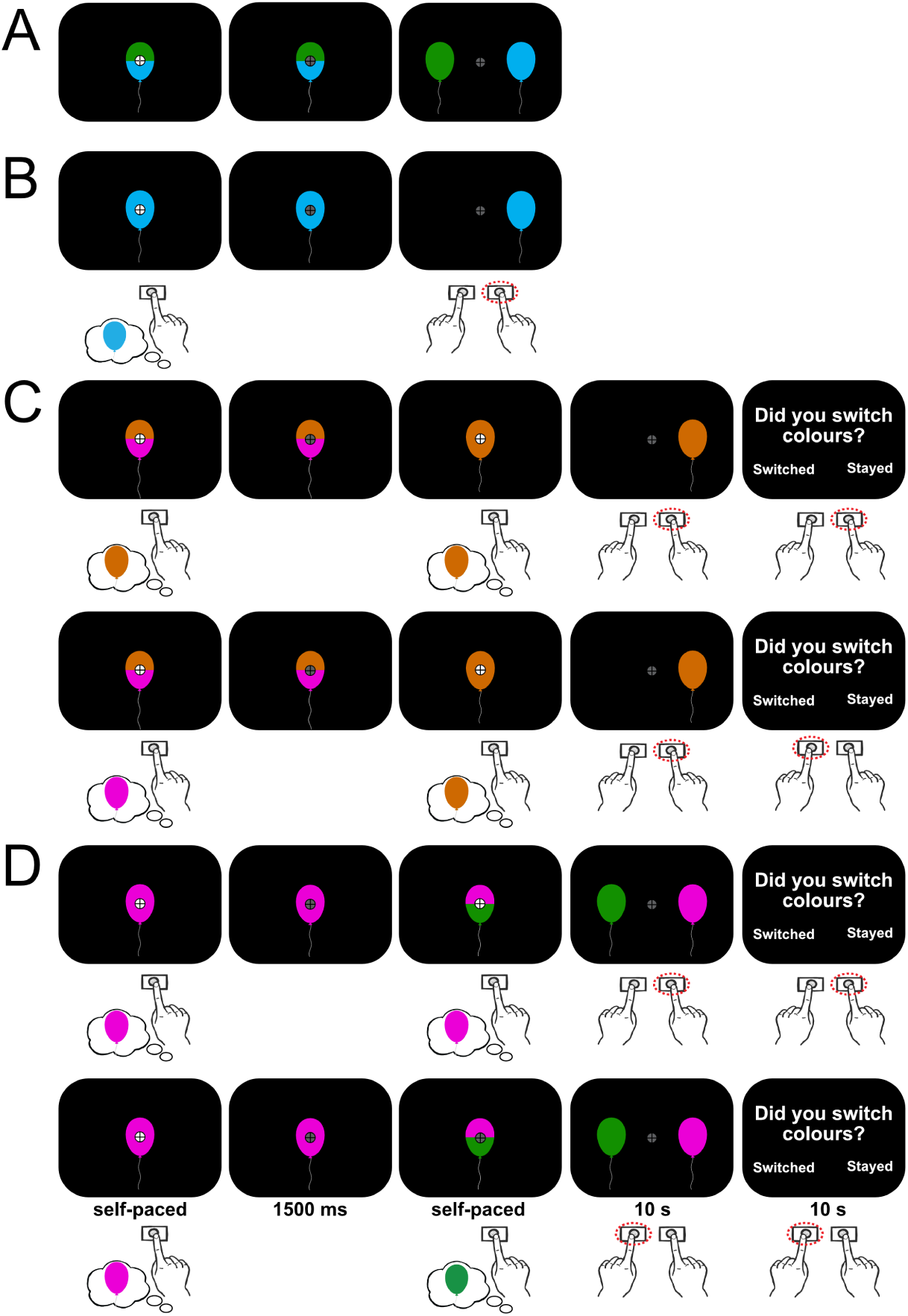
Task design. A) Voluntary decision trials. B) Forced decision trials. In each trial, participants were shown a dual- (voluntary) or single-coloured (forced) balloon and made a keypress with their right middle finger as soon as a decision was made. Balloons of a single colour were then presented at the left and/or right of fixation and participants then indicated the colour they had originally chosen. C) Voluntary-to-forced decision trials. After an initial voluntary decision was made, one of the two colours was removed, forcing participants to either maintain their initial choice or switch. D) Forced-to-voluntary decision trials. After the sole presented option was chosen, a new colour was introduced, allowing participants to maintain or switch their initial choice. In these trials, participants indicated their final choice when balloons appeared to the left and/or right of fixation. They subsequently indicated whether their final choice was the same or different to their initial choice.

### 2.3 Procedure

Participants were seated 50 cm from the monitor in a dark, soundproof room, with their heads stabilised using a chin rest. The experiment included two decision types: voluntary decisions and forced decisions. In voluntary trials (Figure 1A), participants viewed a dual-coloured balloon, indicating two available colour choices. In forced trials, they saw only a single colour (Figure 1B). To encourage more endogenously-driven choice behaviour while keeping voluntary decisions largely free from external constraints (Haggard, 2008), participants were given a cover story to “*collect a colourful range of balloons to decorate a rainbow-themed party*”. This also avoided explicit value-based elements (e.g., rewards or penalties) that could bias strategic or reward-related choice behaviour.

Each trial began with a white fixation stimulus presented for 1200 ms. We instructed participants to maintain a central fixation whenever it appeared. The decision phase then began with colour choices presented on-screen until participants pressed the “+” key on the number pad with their right middle finger to record the moment a decision had been made. Deadlines were not imposed during this decision phase to preserve the voluntary nature of the decisions and to avoid introducing urgency effects. Responses were constrained to the right hand to control for stimulus-response mapping and action selection. Following their response, the fixation stimulus turned grey for 1500 ms and the balloon remained on-screen. Following this, either one or two balloons (corresponding to the colour choices in the trial) were presented to the left and/or right of fixation. Participants pressed keys with their left or right index fingers to indicate the colour they had originally chosen in that trial. Our analyses focused on the initial decision phase, wherein a right-handed "+" keypress indicated decision commitment (i.e., decisional RT). Participants reported which colour they had chosen with a subsequent left/right keypress.

On half of the trials, a second "change-of-mind" (CoM) decision phase followed. This CoM decision phase is not directly relevant to the research questions of the present paper but we describe it for transparency and completeness. For initially voluntary trials, one of the two initial options was removed, forcing participants to either maintain their original choice or switch to the remaining option (Figure 1C). For initially forced trials, a new colour option was introduced, allowing participants to either maintain or switch their initial choice (Figure 1D). Following this CoM decision phase, we presented the available option(s) to the left and right of fixation, and participants indicated their chosen colour using left- or right-labelled keys (i.e., "1" and "3" keys) with their corresponding index fingers. Participants then reported whether they had switched or stayed with their initial decisions. Participants had 10 seconds to report their colour choices and whether they had switched colour choices. CoM trials were not signalled in advance and occurred with fixed probability, so participants made the initial keypress decision under the same expectations across trials; this later report also served as a verification of the initially committed choice.

Participants completed 8 blocks of 36 trials (288 trials in total). Each block lasted approximately 6 minutes, followed by a self-paced break (minimum 10 seconds). Before the main experiment, participants completed one practice block of 36 trials, with the option to attempt up to two additional practice blocks. We report analyses of EEG data from the initial decision phase only. Analyses of EEG data from the CoM decision phase are not reported here but will be the focus of a future publication.

### 2.4 EEG Data Acquisition and Processing

We recorded EEG at a sampling rate of 512 Hz using a BioSemi ActiveTwo system with 64 scalp electrodes, using common mode sense (CMS) and driven right leg (DRL) electrodes (http://biosemi.com/faq/cms/&/drl.htm). We placed an additional eight electrodes on the left and right mastoids, 1 cm from the outer canthi of each eye, and above and below the centre of each eye.

We processed the data using EEGLAB v2023.1 (Delorme & Makeig, 2004) running in Matlab R2022b. First, we identified excessively noisy channels by visual inspection (mean number of bad channels = 0.24, range 0–3) and excluded these from average reference calculations and Independent Component Analysis (ICA). Next, we removed sections with large amplitude artefacts by visual inspection, referenced the data to the average of all the channels, and low-pass filtered the data at 40 Hz. We removed channel AFz to compensate for the rank deficiency introduced by average referencing. We then duplicated the dataset for ICA and high-pass filtered at 0.1 Hz to improve stationarity for ICA. We identified and removed independent components associated with eye blinks and saccades from the unfiltered dataset, high-pass filtered the data at 0.1 Hz, and interpolated AFz and any noisy channels using spherical spline interpolation. With the resulting data, we created stimulus-locked segments of EEG data spanning –2000 to +4000 ms relative to stimulus onset in the initial decision phase and baseline corrected using the 200 ms pre-stimulus interval. Epochs with signals at any scalp channel exceeding ±200 µV were excluded.

In speeded decision-making tasks, EEG signals time-locked to stimulus onset can influence ERP waveforms that are time-locked to the decision and motor response. To minimise the influence of overlapping stimulus-locked activity, we used the RIDE toolbox (Ouyang et al., 2015), following the approach of Steinemann et al. (2018). The RIDE toolbox employs an algorithm that deconvolves stimulus-locked and non-stimulus-locked EEG signals, and has superior performance over comparable methods (Ouyang et al., 2017, 2020). We used the RIDE algorithm to decompose single-trial stimulus-locked data into three subcomponents for each participant using the following time windows: (1) an S (stimulus-locked) subcomponent (0 to 800 ms post-stimulus), (2) an R (response-locked) subcomponent (-600 to -200) ms relative to response onset), and (3) a C (time-varying) subcomponent (200 to 1000 ms post-stimulus) that captures neural activity that differs in latency from trial to trial (e.g., evolving decision formation processes) and not strictly time-locked to either stimulus or response. We included the three subcomponents to improve the accuracy of the S subcomponent estimate in the presence of co-occurring, non stimulus-locked activity that is captured by the C and R subcomponents. Following Sun et al. (2024), we used an extended estimation window for the S subcomponent, relative to default window recommendations in Ouyang et al. (2015), to conservatively estimate and remove stimulus-aligned EEG signals. This addresses concerns noted by Frömer et al. (2024) that response-aligned signals (e.g., evidence accumulation) may artefactually reflect overlapping stimulus-evoked activity. Following best practices implemented in Steinemann et al. (2018) and Sun et al. (2024), we subtracted the estimated S subcomponent (see Supplementary Figure S1A and B) from each stimulus-locked epoch before deriving stimulus- and response-locked segments of EEG data. This subtraction removes estimated stimulus-locked EEG signals, allowing the remaining waveforms to more selectively reflect time-varying and response-locked dynamics (i.e., decision formation and motor preparation). Note that we did not use the R or C subcomponent estimates for our analyses. We also provide plots without ERP deconvolution applied for interested readers in Supplementary Figure S1C–I. Finally, we created response-locked epochs spanning -1200 to +300 ms relative to the response from the resulting S-subcomponent-subtracted stimulus-locked epochs to retain the same pre-stimulus baseline for both stimulus- and response-locked signals (as done in Feuerriegel et al., 2022). This approach allowed us to use a baseline for response-locked epochs that minimises contamination from ongoing motor processes – especially pertinent for voluntary decisions, where the onset of volition is difficult to determine (Haggard, 2008). We then converted the epoched data into current source density (CSD) estimates using the CSD Toolbox with the recommended default toolbox parameters (m-constant = 4 and *λ* = 0.00001; Kayser and Tenke, 2006). These settings match previous studies that applied CSD transforms to examine centro-parietal and motor-related signals (e.g., Corbett et al., 2023; Feuerriegel et al., 2021; Kayser et al., 2007; Pearce et al., 2023). CSD highlights local voltage changes and diminishes the effect of diffuse activity, thereby improving the topographical resolution of EEG signals. This step was crucial for minimising signal overlap between co-occurring signals generated at frontal and central electrodes (e.g., LRP and MB Power) and more posterior electrodes (CPP) around the time of the response (Feuerriegel et al., 2022; Kelly & O’Connell, 2013; O’Connell et al., 2012). After EEG processing, our final response-locked EEG dataset consisted of an average of 258 epochs out of 288 trials retained per participant (range: 214–285). Of the rejected trials, an average of 1 trial per participant (range: 0–8) was excluded due to long RTs (>3700 ms) that caused the response-locked window to extend beyond the specified stimulus epoch window.

### 2.5 Behavioural Analyses

We assessed whether RTs varied between voluntary and forced decisions. Longer voluntary RTs would indicate that participants engaged with the task appropriately, as choosing between multiple options should take longer than single-option, forced-choice decisions. We analysed responses made within 200–3700 ms of the stimulus. For each participant, we computed the mean RTs for voluntary and forced trials and compared these participant-level means using a two-tailed paired-samples t-test. We also performed the same t-tests for the CoM-related RT comparisons and report these results in the Supplementary Material.

### 2.6 EEG Analyses

We analysed response-locked data to assess accumulation-to-bound dynamics corresponding to evidence accumulation and motor preparation before the response. We measured the CPP at electrode Pz, as done in Feuerriegel et al. (2021), Steinemann et al. (2018), and Tagliabue et al. (2019) to index evidence accumulation, and both MB amplitudes and the LHRP at electrode C3 (left motor cortex, contralateral to the right hand) to index motor preparation.

To derive MB time-frequency amplitudes estimates, we applied complex Morlet wavelet convolution to the CSD-transformed data (1–30 Hz in 1 Hz steps, with wavelet cycles increasing linearly from 3 to 10 across the frequency range). Time-frequency amplitude estimates were converted to decibels (dB) relative to the median amplitude across all trials for each participant, using a baseline from -500 ms to -300 ms relative to stimulus onset to minimise contributions from the fixation stimulus presentation. We then created response-locked epochs from the dB-transformed, baseline-corrected data. MB amplitudes were averaged across the 8–30 Hz range (as done in e.g., Feuerriegel et al., 2021; Steinemann et al., 2018; Twomey et al., 2016) at electrode C3.

#### 2.6.1 Characterising Evidence Accumulation and Motor Preparation Signatures of Voluntary Decisions

We tested whether the CPP, MB, and LHRP exhibited hallmark dynamics of evidence accumulation during the initial decision phase: (i) pre-response slopes scaling inversely with RTs (consistent with temporal integration of decision evidence), and (ii) pre-response amplitudes converging across RTs (consistent with a fixed decision threshold). We performed separate analyses for voluntary and forced choice conditions.

We defined pre-response amplitudes as the mean signal from –130 ms to –70 ms relative to the response, capturing immediate pre-response activity while accounting for an estimated 70 ms motor delay between decision and action execution (Kelly et al., 2021), and avoiding earlier amplitude dynamics associated with changes in build-up rates (Feuerriegel et al., 2022). We estimated slopes by fitting a linear regression model to each single-trial response-locked epoch from –300 to –70 ms, in line with prior work (Feuerriegel et al., 2021; Steinemann et al., 2018).

To relate RTs to amplitude and slope measures, we fit linear mixed-effects models (LMMs) with these measures as the outcome and RT as the predictor. As voluntary RTs were slower, more positively skewed, and more variable than forced decision RTs, and because RT scales differed across participants, we log-transformed RTs to reduce the influence of extreme values and to improve the linear approximation. We then z-scored the log-transformed RTs within each participant, removing between participant RT differences so we could focus our inference on within-participant variance (Baayen et al., 2008; Lo & Andrews, 2015). We used a maximal random slopes model that included participant intercepts and slopes for RT, allowing the strength of the RT–EEG measure relationship to vary across participants (Barr, 2013):

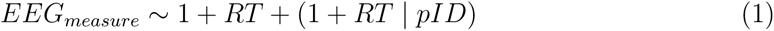

If the random-slopes model produced a singular fit or failed to converge (e.g., due to limited trials, near-zero variance components, or collinearity), we simplified the random-effects structure by removing the random slope and retained a random intercept (Bates et al., 2015):

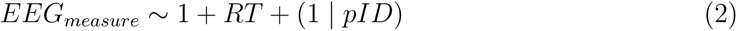

We report regression coefficients (*β* values) from the most complex model that converged without singularity issues. We provide results from both the random-intercept and random-slope models in Supplementary Tables 3–5. For transparency, we also report likelihood-ratio model comparisons using maximum likelihood for the random slope and random intercept models against their respective null models (i.e., without the fixed effect of RT) with matched random effects structures in Supplementary Table 2. Importantly, we did not compare any sets of models that included different random effects structures.

To complement the a priori window-based analyses described above, we also performed mass-univariate analyses across the response-locked epochs (as in Kelly and O’Connell, 2013) with cluster-based permutation corrections for multiple comparisons (Maris & Oostenveld, 2007) implemented in the Decision Decoding Toolbox (Bode et al., 2019). This allowed us to assess whether RT-related effects were localised to the same pre-response interval where CPP–RT effects were reported in the perceptual decision task by Steinemann et al. (2018), while allowing for the possibility that the timing of these effects may differ slightly in voluntary decisions and/or for MB amplitudes and the LHRP, but still remain proximal to the response. For each trial and time point, we computed CPP, MB, and LHRP amplitudes and estimated linear slopes in 300 ms windows (as per minimum window width recommendations from Cohen, 2019 for effective temporal resolution of Morlet MB estimates) centred on each time point. For slope estimates, we excluded the first and last 150 ms where a full 300 ms window could not be created.

We then fit regression models to predict slope and amplitude values at each time point using RT as a predictor. One-sample t-tests were performed to compare the distributions of the resulting *β* values against zero. We formed clusters from neighbouring time points with *p < .*01 (uncorrected), and summed *t*-values to compute cluster masses. We permuted condition labels for a subset of participants (10,000 permutation iterations) to build an empirical null distribution using the largest observed cluster mass per permutation. Clusters exceeding the 97.5*^th^* percentile were deemed significant (*α* = .05, two-tailed), controlling for the weak family-wise error rate (Groppe et al., 2011; Maris & Oostenveld, 2007). Surviving clusters denoted time windows where amplitudes and slopes covaried with RT.

Using the BayesFactor toolbox v3.0 (Krekelberg, 2024), we also derived Bayes Factors (BF_10_) from Bayesian one-sample *t*-tests with default Cauchy priors (r = 0.707; Rouder et al., 2017). The BF_10_ is a ratio that reflects the likelihood of observing the data under the alternative hypothesis as compared to the null. We interpret BF_10_ as follows: BF_10_ > 3 positive, BF_10_ > 20 strong, and BF_10_ > 150 very strong preferential evidence for the alternative hypothesis (Kass & Raftery, 1995). Please note that Bayes factors were computed separately from the cluster-based permutation tests and were not used to determine significance for the multiple-comparisons correction. Bayes factors are reported here solely as a descriptive index showing where evidence supports the presence versus absence of an effect at each time point.

Finally, we compared group-averaged waveform amplitudes of the three signals between voluntary and forced decisions using mass-univariate paired-samples *t*-tests, using the same cluster-based permutation correction as above. This allowed us to determine whether voluntary and forced decisions produced any systematic differences in the morphologies of each signal.

## 3 Results

To validate our task manipulation, we tested whether RTs differed between voluntary and forced decisions. Individual and group mean RTs for each decision type are plotted in Figure 2A. Initial forced decisions (*M* = 0.86 s) were faster than voluntary decisions (*M* = 1.10 s), *t*(48) = 6.07, *p < .*001, 95% CI [0.15, 0.29], Cohen’s *d_z_* = 0.87). RT distributions pooled across participants shown in Figure 2B were positively skewed for both voluntary and forced decisions, consistent with the ex-Gaussian-like distributions typically observed in decision tasks. Individual participant RT distributions are displayed in Supplementary Figure S2. Crucially, forced decision RTs were much slower than typical simple reaction time (∼ 235 ms; Woods et al., 2015) and discrimination time (∼ 150 ms; Hopf et al., 2002) latencies, supporting the interpretation that forced decisions entailed colour identification and decision commitment rather than reflexive detection.

**Figure 2.**
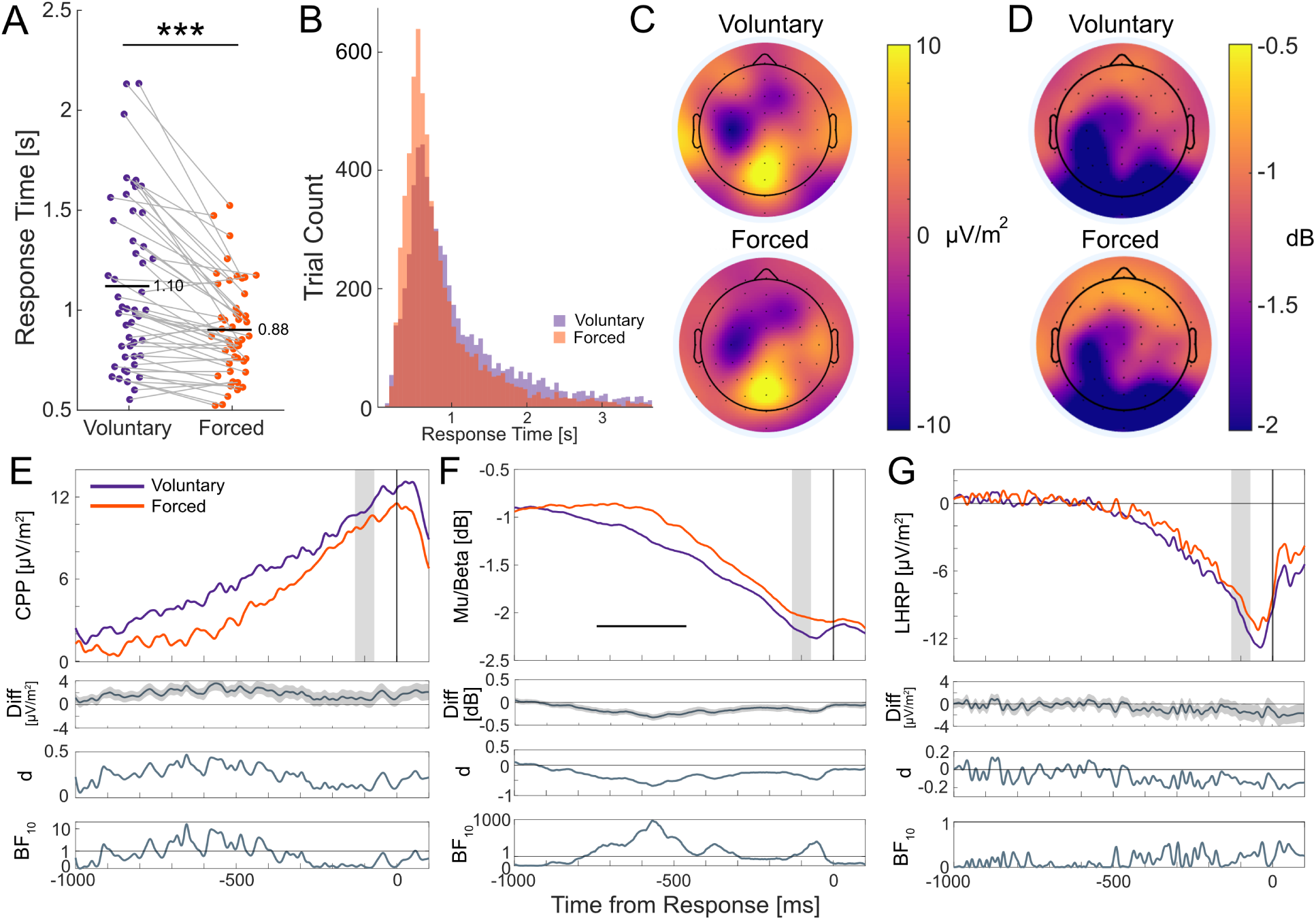
Mean response times across decisions and EEG signal dynamics. A) Participant (dots) and group (horizontal black lines) mean response times for voluntary and forced decisions. Grey lines connect paired observations from the same participant. Asterisks denote statistically significant response time differences (p < .001). B) Pooled response time distributions for voluntary (purple) and forced (decisions), depicting positively skewed distributions across both decision types. C - D) Response-locked scalp topographies of voluntary (top) and forced (bottom) decisions averaged across the -130 to -70 ms window preceding response onset. Panel B shows time-domain ERP amplitudes; Panel C shows Mu/Beta (8–30 Hz) spectral amplitudes. E) Group-averaged CPP, F) MB amplitudes, and G) LHRP for voluntary (purple) and forced (orange) waveforms with accompanying difference waves (and shaded standard error regions), effect size estimates (Cohen, 1988), and BFs in favour of the alternative hypothesis (i.e., amplitude differences) at each time point. Grey areas represent the -130 to -70 ms window used to derive the scalp maps. The horizontal black line under MB waveforms denotes statistically significant differences after correcting for multiple comparisons. Horizontal lines in BF plots denote BF_10_ = 1, indicating a lack of preferential support for either the alternative or null hypothesis. Overall, all three signals exhibited the expected ramping morphology, with no reliable amplitude differences between voluntary and forced decisions immediately before the response.

To provide additional insight into choice behaviour for voluntary decision trials, we report each participant’s colour choice frequencies and mean RTs for each colour in Supplementary Table 1, alongside post-experiment self-reported preference rankings and decision strategies. These summaries show patterns consistent with preference- and goal-driven choice behaviour for many participants, while also highlighting some between-participant heterogeneity in choice behaviour. Crucially, these patterns suggest that participants engaged with voluntary trials by applying endogenously motivated decision strategies.

### 3.1 Characteristics of the CPP, MB, and LHRP Signals

Pre-response amplitude scalp maps for both voluntary and forced decisions in Figure 2C showed a prominent centro-parietal positivity focused at Pz, which was spatially distinct from motor-related activity over the left motor cortex at C3. MB scalp topographies in Figure 2D also showed focal power reduction (i.e., more negative dB values) over left-central sites near C3 for both decision types. Importantly, in this CSD-transformed data, a typical RP-like negativity focused over Cz was not apparent in this single-effector response design. Across voluntary and forced decisions, all three neural signals exhibited the expected ramping morphology leading up to the motor response: a positive-going CPP build-up (Figure 2E) and negative-going build-ups for MB amplitudes (Figure 2F) and the LHRP (Figure 2G).

To test for any clear differences across decision types, we compared voluntary vs forced waveform amplitudes using mass-univariate analyses. Across all three signals, we did not find reliable waveform differences within time windows immediately prior to the response. For MB amplitudes, an early significant cluster emerged between -740 and -460 ms (cluster p < .001). This window overlapped with a period where CPP BFs indicated moderate support for amplitude differences but this did not survive multiple testing corrections. These early effects likely reflect ramp-onset timing differences driven by RT differences between voluntary and forced decisions, as shown in Figure 2A, and lay outside our pre-response window of interest. We also found moderate BFs in support of MB amplitude differences around response onset (-100 to -10 ms), but these effects also did not survive correction for multiple comparisons. The LHRP did not show statistically significant amplitude differences at any time point. Given these canonical profiles, we proceeded to test accumulation-to-bound dynamics across the three signals.

### 3.2 The CPP exhibits accumulation-to-bound dynamics in voluntary decisions

We examined whether the CPP, MB amplitudes, and the LHRP exhibited accumulation-to-bound characteristics: (i) build-up slopes that scaled inversely with RTs, and (ii) pre-response amplitudes that converged to similar levels irrespective of RTs. These dynamics were assessed in two ways: first, using fixed pre-response windows for slopes (-300 to -70 ms) and amplitudes (-130 to -70 ms) as defined in prior perceptual decision-making work (e.g., Feuerriegel et al., 2021; O’Connell et al., 2012; Steinemann et al., 2018); second, using mass-univariate analyses across the full response-locked epoch to identify any additional RT-related effects outside the pre-defined windows. Here, we present data relative to the initial decision in each trial. Data relating to CoM decisions are beyond the scope of our paper.

To visualise these results, Figures 3A and B show group-averaged stimulus-locked (left) and response-locked (right) waveforms for voluntary and forced decisions, respectively, split by RT bins (fast, medium, and slow). Trials were split into three equal-sized RT bins for each participant and averaged across participants. Response-locked waveforms in Figures 3A and B (right panel) show the expected gradual build-up across RT bins – with steeper build-ups for the fastest RT bin – and amplitudes converging to similar pre-response levels. We also include stimulus-locked waveforms at Pz in Figures 3A and B (left panel) for completeness and descriptive context, but we do not analyse them further here. In our task, the visual display changed immediately after a keypress response. Therefore, stimulus-locked waveforms capture a combination of pre- and post-decisional ERPs, including visual evoked responses, whose timing varies with RT. Accordingly, stimulus-locked plots should be interpreted descriptively and not as a direct readout of decision dynamics.

**Figure 3.**
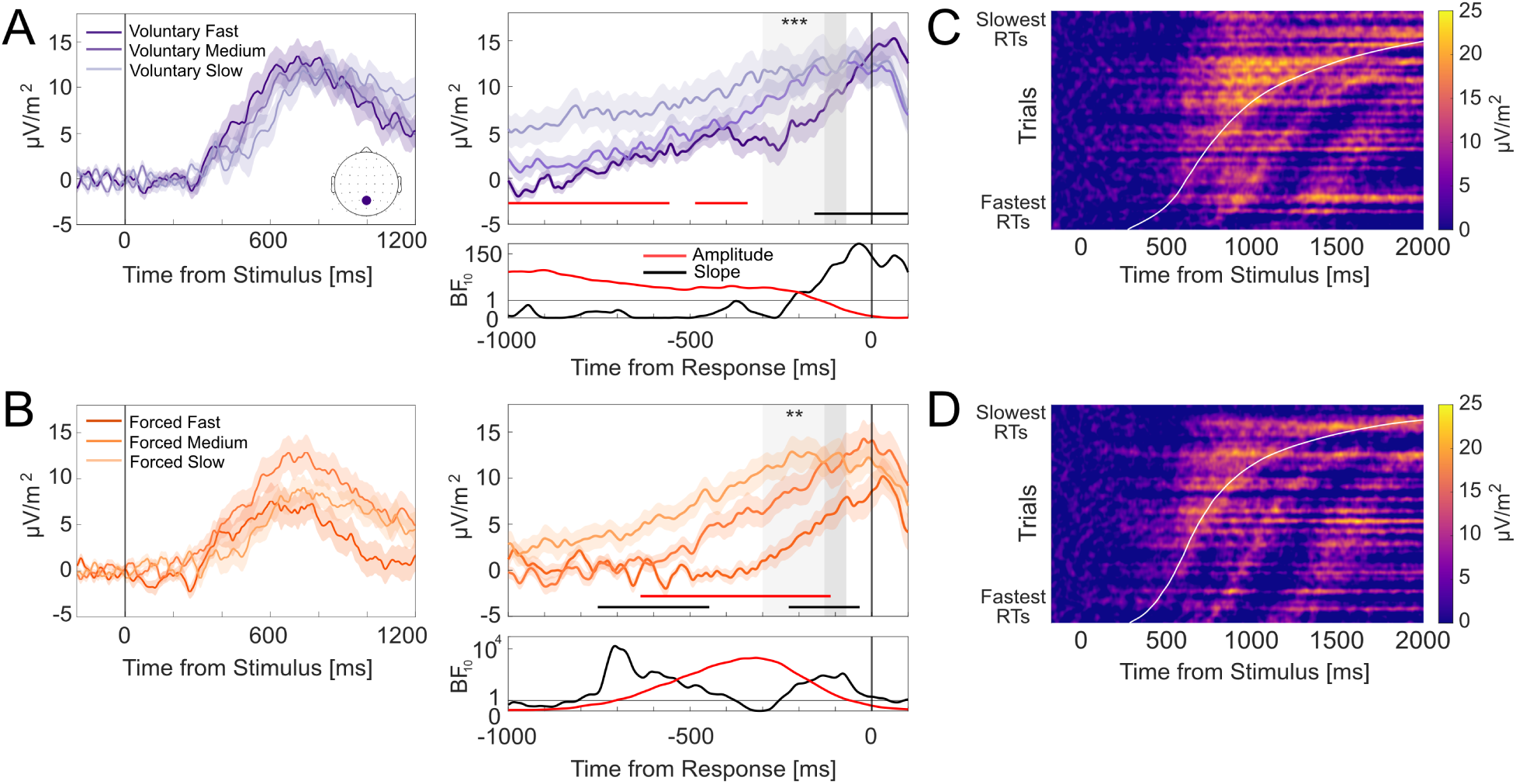
CPP evidence accumulation dynamics at electrode Pz. A-B) Group-averaged stimulus-locked (left) and response-locked (right) ERP waveforms for voluntary (A) and forced (B) decisions split by fast, medium, and slow RT bins, with shading denoting standard errors. Dark grey areas denote the -130 to -70 ms pre-response amplitude window. Light grey areas denote the -300 to -70 ms time slope window. Asterisks denote statistically significant RT associations with ERP amplitudes and slope differences within their respective windows (** denotes p < .01; *** denotes p < .001). Red and black horizontal lines denote windows where RTs were associated with ERP amplitudes and slope differences, respectively. Corresponding BF_10_ estimates for each association are plotted below. Horizontal lines in BF plots denote BF_10_ = 1, indicating a lack of preferential support for either the alternative or null hypothesis. Bayes factors for forced decisions are plotted on logarithmic scales due to the wide ranges of values across the time-course. C-D) Single-trial stimulus-locked CPP amplitudes for voluntary (C) and forced (D) decisions pooled across participants. Trials are sorted by RT (fastest below and slowest on top). Data were smoothed over 300-trial bins using a sliding Gaussian-weighted moving average filter. RTs are denoted by the curved white line.

Within the pre-defined analysis windows, faster RTs were associated with steeper CPP slopes for both voluntary (*β* = −0.02, *SE* = 0.005, *t*(48.29) = −4.03, *p < .*001) and forced (*β* = −0.02, *SE* = 0.01, *t*(41.81) = −3.34, *p* = .002) decisions. Negative coefficients indicate steeper build-up rates for faster RTs and shallower slopes for slower RTs.

Mass-univariate analyses also revealed RT effects that overlapped with these pre-defined windows. Voluntary RTs covaried with CPP slopes from approximately -160 to +115 ms (cluster p < .001), and forced RTs covaried from –230 to -30 ms (cluster p = .002) and at an earlier window spanning –900 to -300 ms (cluster p < .001). BFs indicated very strong support for RT-slope associations during these windows. Together, these findings indicate that CPP slopes scale systematically with RTs in both voluntary and forced decisions, consistent with the CPP indexing an evolving decision variable.

CPP pre-response amplitudes did not vary systematically with RTs for both voluntary (*β* = 1.55, *SE* = 1.34, *t*(50.38) = 1.15, *p* = .255) and forced (*β* = 1.20, *SE* = 0.75, *t*(6173.33) = 1.60, *p* = .109) decisions. Additionally, mass-univariate analyses

Figure 3 only identified RT-amplitude associations for voluntary decisions, spanning -1050 to –555 ms (cluster p = .013) and -485 to -340 ms relative to the response (cluster p < .05); and for forced decisions from –635 to –115 ms (cluster p = .004). These effects did not extend into the time period just preceding response onset, further suggesting that CPP amplitudes converged to similar levels around the time of the motor response.

Taken together, CPP slopes scaled reliably with RTs, while pre-response amplitudes converged to similar levels irrespective of RTs. To further visualise these accumulation-to-bound dynamics, we plotted heatmaps of single-trial stimulus-locked CPP amplitudes at electrode Pz, sorted by ascending RTs and pooled across participants for voluntary (Figure 3C) and forced decisions (Figure 3D; as done in O’Connell et al., 2012 and Feuerriegel et al., 2022). These heatmaps provide an intuitive representation of tracing how amplitude build-up unfolds as a function of RTs, reflecting temporal integration of evidence to a decision bound. For both voluntary and forced decisions, the heatmaps show a gradual positive-going build-up in CPP amplitudes (i.e., increasing yellow saturation) at similar onsets across trials that culminate at comparable peak levels around the time of the response (denoted by the curved white line).

### 3.3 MB amplitudes exhibit accumulation-to-bound dynamics in voluntary decisions

Response-locked MB amplitude trajectories for voluntary and forced decisions, split by fast, medium, and slow RT bins, are displayed in Figures 4A and B, respectively. Both figures show MB amplitudes generally converged to a common value prior to the response and the rate of amplitude decrease was more gradual for slower RTs. Faster RTs were associated with steeper MB slopes for voluntary decisions (*β* = 0.002, *SE* = 0.001, *t*(5619.08) = 1.47, *p < .*001) but not for forced decisions (*β* = 0.0001, *SE* = 0.001, *t*(6054.89) = 0.11, *p* = .141) decisions. However, outside of this window, mass-univariate analyses detected significant RT effects spanning -450 to -345 ms for voluntary decisions (cluster p = .039), and between -915 and -730 ms (cluster p = .004) and -365 to -50 ms (cluster p < .001) for forced decisions. This indicates that for both voluntary and forced decisions, faster RTs were accompanied by steeper (i.e., more negative) MB slopes, whereas longer RTs showed shallower (i.e., less negative) slopes.

**Figure 4.**
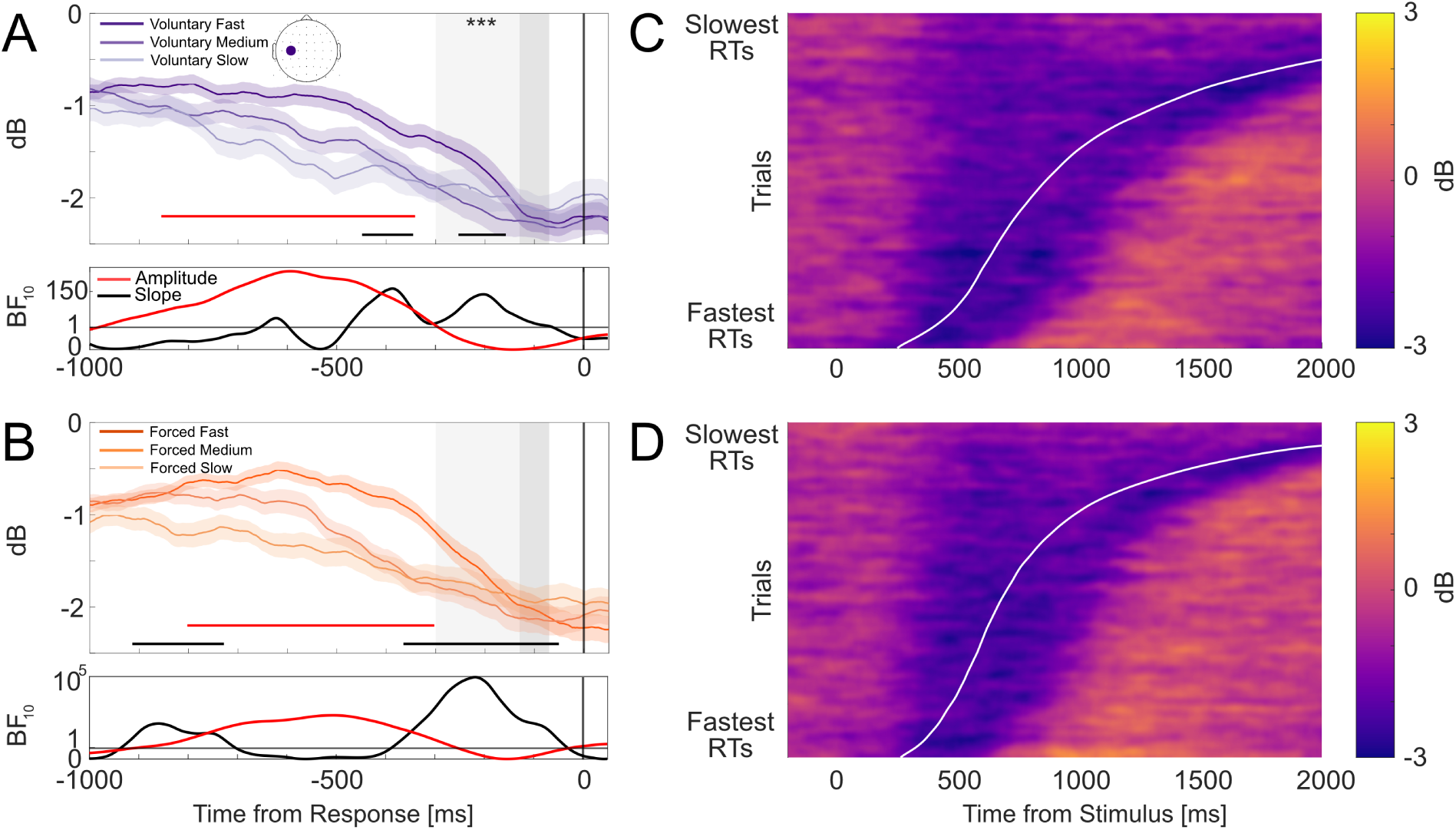
Mu/Beta (8–30 Hz) dynamics at C3 (scalp map in top-left panel). A-B) Group-averaged response-locked waveforms for voluntary (A) and forced (B) decisions split into fast, medium, and slow RT bins, with shading denoting standard errors. Dark grey areas denote the -130 to -70 ms pre-response amplitude window. Light grey areas indicate the -300 to -70 ms slope window. Asterisks denote statistically significant RT-slope associations for the fixed window analysis (*** denotes p < .001). Red and black horizontal lines denote windows where RTs were associated with amplitude and slope differences, respectively. Corresponding BF_10_ estimates for each association are plotted below. Horizontal lines in BF plots denote BF_10_ = 1, indicating a lack of preferential support for either the alternative or null hypothesis. Note that for (B), BFs are plotted on logarithmic scales due to the wide ranges of values across the time-course. C-D) Heat maps depicting MB amplitudes for voluntary (C) and forced (D) decisions relative to stimulus onset. Trials are sorted by RT (fastest below and slowest on top). Data were smoothed over 300-trial bins using a sliding Gaussian-weighted moving average filter. RTs are denoted by the curved white line.

MB pre-response amplitudes did not vary systematically with RTs for both voluntary (*β* = 0.03, *SE* = 0.07, *t*(54.46) = 0.43, *p* = .665) and forced (*β* = −0.06, *SE* = 0.09, *t*(47.34) = −0.75, *p* = .456) decisions. Similarly, mass univariate analyses did not reveal support for RT-amplitude effects, particularly in the 400 ms interval before the response.

Heatmaps of single-trial MB amplitudes for voluntary (Figure 4C) and forced (Figure 4D) decisions both show accumulation-to-bound dynamics closely resembling those of the CPP: gradual decrease in amplitudes, indexed by increasingly purple saturation, beginning at comparable onsets across trials and peaking at similar amplitude levels around the response.

### 3.4 The LHRP exhibits threshold-crossing dynamics but may not trace evidence accumulation

Figures 5A and B show response-locked group-averaged LHRP waveforms for voluntary and forced decisions, respectively, split into fast, medium, and slow RT bins. Both figures show LHRP waveforms that gradually became more negative-going and reached their peak just prior to the response. Although forced decision waveforms in Figure 5B display visual separation across RT bins (like in the CPP), these separations are likely accentuated by the highly skewed RT distributions. These waveforms should be interpreted with the same caution noted for the CPP.

**Figure 5.**
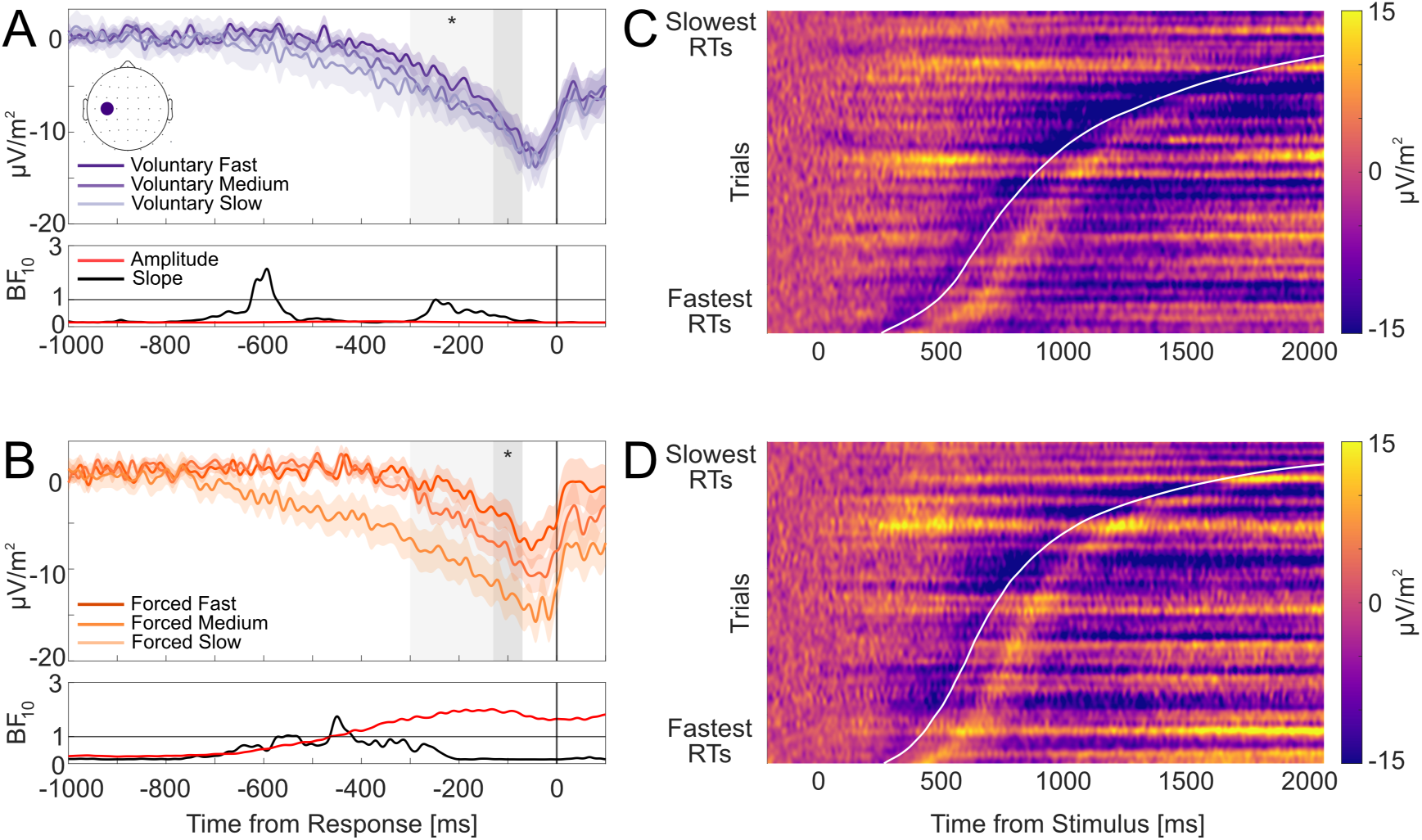
Left hemisphere readiness potentials at electrode C3 (scalp map in top-left panel). A-B) Group-averaged response-locked waveforms for voluntary (A) and forced (B) decisions split into fast, medium, and slow RT bins, with shading denoting standard errors. Dark grey areas denote the -130 to -70 time pre-response amplitude window. Light grey areas indicate the -300 to -70 ms slope window. Asterisks denote statistically significant RT associations between ERP amplitudes and slope differences within their respective windows (* denotes p < .05). Corresponding BF_10_ estimates in red and black denote preferential support for RT associations with amplitude and slope differences, respectively, at each time point are plotted below. Horizontal lines in BF plots denote BF_10_ = 1, indicating a lack of preferential support for either the alternative or null hypothesis. However, we did not observe any significant clusters in which RT modulations on the amplitudes and slopes were sustained over time. C-D) Heat maps depicting LHRP amplitudes for voluntary (C) and forced (D) decisions relative to stimulus onset. Trials are sorted by RT (fastest below and slowest on top). Data were smoothed over 300-trial bins using a sliding Gaussian-weighted moving average filter. RTs are denoted by the curved white line.

Within the pre-defined analysis window, RTs significantly predicted LHRP slopes for voluntary decisions (*β* = 0.01, *SE* = 0.005, *t*(49.9) = 2.42, *p* = .019), indicating that slower RTs were associated with shallower LHRP slopes. However, mass-univariate analyses did not reveal any reliable RT effects at any time point. As cluster-based permutation tests are more sensitive to detecting effects that span across several consecutive time points, the absence of significant clusters found suggests that RT effects found during the pre-defined window may instead reflect local variability or effects limited to that isolated time window. Furthermore, BFs across time points within the pre-defined window were less than 1, indicating a lack of clear evidence for RT-slope effects. For forced decisions, RTs did not significantly predict LHRP slopes (*β* = −0.01, *SE* = 0.01, *t*(48.48) = −1.03, *p* = .306). Mass univariate analyses and BFs also did not provide substantial evidence in support of reliable RT-slope effects at any time point.

RTs did not significantly predict LHRP pre-response amplitudes for voluntary decisions (*β* = −0.56, *SE* = 0.99, *t*(6038) = −0.57, *p* = .571). In contrast, for forced decisions, RT was a significant predictor of amplitudes (*β* = −3.53, *SE* = 1.34, *t*(45.37) = −2.62, *p* = .012), with slower RTs associated with more negative-going amplitudes. However, mass-univariate analyses and BFs did not provide preferential support for RT-amplitude associations at any time point for voluntary decisions nor substantial support for forced decisions (*BF*_10_ < 3), which may similarly indicate that effects during the pre-defined time window were not sustained for long enough beyond the localised time window or sufficiently strong to survive corrections for multiple comparisons.

The heatmaps for voluntary (Figure 5C) and forced (Figure 5D) decisions both show negative-going build-up that peaks at similar amplitude levels around response onset regardless of RT. However, the LHRP build-up begins at a relatively fixed time before the response in contrast to earlier build-ups observed in the CPP and MB, reflecting dynamics that align with a late-stage, effector-specific motor preparation rather than extended evidence integration.

Taken together, the analyses do not yield strong support for RT modulation of LHRP slopes or pre-response amplitudes. However, the visual patterns in Figure 5B and localised effects in the pre-defined slope and amplitude windows leave open some possibility for subtle RT-related influences that may have been too weak to detect in our data.

## 4 Discussion

We assessed whether neural signatures of evidence accumulation and motor preparation in perceptual decision tasks also characterise voluntary decisions, which we operationalised in our experiment as endogenously guided, two-alternative choice decisions, which are primarily driven by abstract goals and preferences. We also examined forced decisions whereby a single colour specified the response, similar to perceptual decision-making designs that require stimulus evaluation and response commitment, even though choices were externally specified (Forstmann et al., 2016; Gold and Shadlen, 2007; such as colour identification tasks). Importantly, our experimental design allowed us to investigate the formation of voluntary decisions independently from action selection or effector choice. which still require stimulus evaluation and response commitment even though choices were externally specified (Forstmann et al., 2016; Gold & Shadlen, 2007).

After accounting for stimulus-locked contributions to response-aligned neural activity, waveform amplitudes and pre-response scalp topographies of CPP, MB and LHRP signals were highly similar between voluntary and forced decisions. Furthermore, both CPP and MB signals displayed accumulation-to-bound properties: (i) build-up rates (slopes) were steeper for faster RTs and (ii) pre-response amplitudes converged to similar levels around the time of the motor response. The LHRP findings were more mixed: we observed an RT–slope association for voluntary decisions and an RT-amplitude association for forced decisions within their respective pre-response measurement windows. However, these effects were not clearly apparent in our mass-univariate analyses and did not survive correction for multiple comparisons. Therefore, our results suggest that the LHRP ramps towards a stereotyped amplitude level before a motor response, but may not trace evidence accumulation in a similar manner to CPP and MB signals. Taken together, these findings point to shared evidence accumulation and motor preparatory mechanisms that underlie internally and externally guided choices.

### 4.1 The CPP traces evidence accumulation during both perceptual and internally-driven voluntary decisions

Recent work by Frömer et al. (2024) has questioned whether the CPP exclusively indexes evidence accumulation in perceptual decision-making. However, O’Connell et al. (2025) posit that true CPP-like dynamics in Fromer and colleagues’ results have been masked by their methodological and analysis decisions, including their choice of deconvolution approach to reduce component overlap. We mitigated these issues by applying an alternative deconvolution approach (i.e., RIDE; Ouyang et al., 2015) to reduce stimulus-related overlap in our response-locked data, and by using CSD transformations to better isolate measures of the CPP from other, co-occurring EEG signals (Feuerriegel et al., 2022; Kelly and O’Connell, 2013). When accounting for these confounds, we observed canonical accumulation-to-bound CPP dynamics in voluntary decisions that could be understood as examples of simple value-based decisions – deliberating between two alternatives based on endogenous factors, including preferences and goals, without the constraints of an objectively “correct” answer. Therefore, in contrary to Frömer et al. (2024), our findings support the view that the CPP could be a neural correlate of a domain-general evidence accumulation process that is not exclusive to perceptual decisions. We also present, to our knowledge, the first evidence of CPP and MB correlates of evidence accumulation in a task commonly used to study voluntary (i.e., intentional) decisions.

Despite qualitative differences in task demands, CPP waveform amplitudes did not differ reliably between voluntary and forced decisions, and both showed accumulation-to-bound properties. In our task, voluntary decisions required recognising that a choice existed and then deliberating between two alternatives based on more endogenously-driven factors, including preferences, which may be stable or fluctuate from moment-to-moment, goals and other internal heuristics) rather than external cues. By contrast, forced decisions followed a fixed rule – choose the sole presented option – but nonetheless entailed a perceptual decision to identify and commit to the colour presented. Crucially, forced decision RTs were slower than typical simple reaction or discrimination task RTs (∼150 - 235 ms; Hopf et al., 2002; Woods et al., 2015), indicating decisional processing rather than reflexive responding. A deliberate decision was required as each colour choice needed to be maintained in memory in anticipation of a possible change of mind in half of the trials. Together, these observations suggest that the CPP tracks a common decision variable irrespective of whether evidence is exogenous or endogenously generated, supporting sequential-sampling accounts of voluntary choice.

### 4.2 MB amplitudes trace the accumulation of decision evidence in the motor system

Like the CPP, MB amplitudes reflected an RT-dependent build-up to a fixed pre-response level for both voluntary and forced decisions, suggesting similar motor-gating policies in which actions are triggered once motor preparatory activity reaches a stereotyped threshold (Donner et al., 2009; Lange et al., 2013; O’Connell et al., 2012). We note that MB trajectories can be sensitive to urgency effects under speeded response deadlines. MB amplitudes can progress toward the response-triggering threshold at a faster rate than expected based on evidence accumulation rates, and can even begin ramping prior to the onset of a decision-relevant stimulus (Kelly et al., 2021; Steinemann et al., 2018). However, our MB results are more consistent with a decision variable within the motor system where there is little speed pressure: when evidence was accumulated slowly, MB ramped more gradually and resembled the trajectories of the CPP. Variations in response onset may better be explained by variations in evidence accumulation rates rather than idiosyncratic changes in motor-execution thresholds (Gold & Shadlen, 2000; Kim & Shadlen, 1999). Furthermore, our task did not explicitly impose speed pressure manipulations – a context in which MB and CPP dynamics are often similar (e.g., Donner et al., 2009; Feuerriegel et al., 2021; Lange et al., 2013; O’Connell et al., 2012).

Our MB dynamics also align with accumulator modelling results in speeded perceptual decision tasks, which reliably show fixed boundary setting with variability in stimulus-driven (e.g., ease of discrimination) or goal-driven (e.g., reward rate) evidence accumulation rates (Brown & Heathcote, 2008; Ratcliff & McKoon, 2008; Smith & Ratcliff, 2004). These dynamics possibly reflect the lack of urgency manipulations and do not support the idea that urgency is an essential feature of voluntary decisions as previously proposed by Brass et al. (2019). Future work could introduce urgency manipulations (e.g., deadlines) to test whether neural signatures of urgency – reduced CPP and stable MB pre-response amplitudes for slower decisions – are observed (Feuerriegel et al., 2021; Kelly et al., 2021; Steinemann et al., 2018).

Having observed CPP and MB dynamics in both voluntary and forced decisions that were consistent with model-motivated accumulation-to-bound predictions, with broadly similar waveform amplitudes across both decisions, our findings suggest that sequential sampling models may extend beyond classical perceptual decision-making tasks to voluntary, two-alternative decisions. Because our inferences of evidence accumulation lies primarily on observing model-predicted dynamics in our EEG data, our inferences can be further supported by future joint modelling (e.g., hierarchical DDM and race models) of voluntary decisions (as done in Kelly et al., 2021 for perceptual decisions and Sun et al., 2025), which provide a more direct test of how CPP and MB dynamics relate to drift and boundary parameters. For example, CPP slopes can serve as a proxy for trial-wise drift rates to link variability in RTs to endogenous evidence, such as individual choice preferences, and CPP pre-response amplitudes could map onto decision-bound criteria. MB signals can be modelled similarly in voluntary decisions made under speed pressure to assess whether MB signals reflect urgency-related boundary shifts – indexed by flexible decision bound criterion and RT-amplitude associations – and whether the CPP consistently tracks evidence accumulation trajectories, as in perceptual tasks (Kelly et al., 2021; Steinemann et al., 2018). Such modelling efforts will advance models of voluntary decisions beyond “when to act” (e.g., Schurger, 2018; Schurger et al., 2012) to “which option” decision dynamics, and further clarify the functional commonalities that underlie evidence accumulation in voluntary and forced decisions.

### 4.3 The LHRP indexes late motor preparation but may not reflect evidence accumulation dynamics

Relative to the CPP and MB amplitudes, the LHRP showed less consistent evidence for accumulation-to-bound trajectories for both voluntary and forced decisions. We observed subtle RT-slope associations in the pre-response window for voluntary decisions. Conversely, in forced decisions, slower RTs were associated with more negative LHRP amplitudes in the –130 to –70 ms window, but this effect was not observed for voluntary decisions. Although this RT-amplitude association could be interpreted as increasing boundaries with elapsed time, the directionality of this pattern in our results runs counter to more commonly-proposed collapsing-bound predictions (e.g., COINTOB model; Brass et al., 2019), where slower responses terminate at lower thresholds (O’Connell & Kelly, 2021; Steinemann et al., 2018). Moreover, these effects were not statistically significant in the mass-univariate analyses after correcting for multiple comparisons.

Although accumulation-like trajectories have previously been reported for the LRP using analyses similar to our study (Kelly & O’Connell, 2013), and for the RP using computational modelling (Lui et al., 2021; Maoz et al., 2019), these measures differ from our LHRP in critical ways that limit their direct comparison. The classical RP is measured at Cz, indexing broader central motor preparatory activity, thought to originate from pre-SMA and the SMA. However, without CSD transformations, activity at Cz can capture overlapping signals from neighbouring sites, such as motor-independent CPP signals from Pz. The LRP is computed as the contralateral–ipsilateral difference between C3 and C4, indexing bilateral effector competition. LRP ramping trajectories arise from participants choosing between hands, yielding a growing lateralised difference in motor preparation.

Meanwhile, in our single-effector design, we mapped every decision to a right-handed keypress, thereby eliminating effector competition and motivating a focus on contralateral activity at C3. This choice is further supported by the scalp maps in Figure 2B, which also did not show an RP at Cz in our CSD-transformed data. Therefore, it is plausible that previously reported evidence accumulation dynamics may in part arise from activity related to action selection.

While we cannot conclusively rule out accumulation-to-bound dynamics here, our results suggest that when action selection is controlled for, the LHRP at C3 may be better interpreted as a late-stage effector-specific gate that releases an action once a relatively fixed motor threshold is crossed. It appears less clearly as an evolving decision variable for the motor system.

### 4.4 Limitations and future directions

Although our single-effector design strengthened our inferences about decision formation independent of hand selection, it also limits direct comparisons with existing RP and LRP findings that show accumulation-like dynamics in dual-handed task designs with effector competition (e.g., Kelly and O’Connell, 2013; Maoz et al., 2019), and in single-effector designs using, for example, shoulder rotation responses in different directions (Lui et al., 2021). Future studies could reintroduce effector competition using dual-handed response designs to test whether the RP or LRP signals (guided by CSD-transformed scalp topographies) show accumulation-to-bound dynamics for voluntary decisions using the same analysis framework. Such analyses will clarify whether readiness potentials exhibit accumulation-to-bound properties only when effector competition is present, or whether they primarily reflect late-stage threshold-crossing properties – akin to our LHRP findings.

There is also the possibility that a subsequent CoM phase could shape participants’ overall response policy during the initial decision phase (e.g., adopting a more cautious commitment threshold, or treating initial choices as more provisional). However, CoM trials were not cued at the start of each trial and occurred with a fixed probability across the experiment. Regardless, participants would still need to make a decision and our RT distributions and choice behaviour data show strong support for genuine decision-making. Therefore, although the CoM manipulation could potentially influence decision strategies or response policies, these are likely to apply across all trials and are unlikely to account for the specific accumulation-consistent signatures we report.

In our study, inferences relating to stimulus-locked ERPs were constrained by our task design. Visual display changes occurred immediately after the keypress response (e.g., fixation stimulus changes), and there were substantial differences in the degree of RT variability across the RT tertile bins. Consequently, plotted waveform amplitudes reflected different mixtures of pre-decisional build-up and post-decisional visual evoked activity as well as different degrees of temporal smearing. Additionally, stimulus-locked ERP slope estimates, using trial-varying windows that are adjusted to end at the RT, can produce spurious slope-RT associations even in the absence of true ramping signals. For these reasons, stimulus-aligned CPP plots and analyses are not straightforward to interpret. A design that delays post-response visual changes would permit cleaner isolation of pre-decisional stimulus-locked activity in future work (e.g., Sun et al., 2025). Nonetheless, our core inferences here were based on response-locked analyses that directly traced dynamics around decision commitment and are not compromised by these issues.

In addition to a prominent CPP and left-hemispheric motor activity, our scalp maps also showed a fronto-central negativity consistent with the contingent negative variation (CNV) ERP component (Figure 2C). The CNV has been proposed to index a movement-independent urgency signal that grows with elapsed time and can reset when actions are withheld, placing this temporally between evidence accumulation and motor execution (McCone et al., 2025). These properties suggest that the CNV may be more tightly linked to the termination of decisional evidence accumulation (e.g., through urgency parameters or collapsing bounds), rather than motor execution. At the same time, CNV trajectories closely resemble the RP in morphology and topography, although their distinct functional roles in decision-making remain unclear (see Schurger et al., 2021 for detailed discussion). We did not focus on the CNV here as our a priori questions concerned decisional evidence accumulation (i.e., CPP) and motor preparation (i.e., MB and LHRP) trajectories. Additionally, our task did not impose explicit urgency manipulations – conditions under which CNV dynamics have been most informative. Future studies could introduce deadlines to test whether the CNV unfolds as an evolving decision variable with accumulation-to-bound trajectories like the urgency-insensitive CPP, or as a time-dependent urgency signal that lowers effective decision thresholds as time elapses (as found in McCone et al., 2025). Observing such CNV–CPP coupling under time pressure provides a decisive test of urgency-based accounts and clarifies how urgency shapes the interplay between decision termination and motor gating in voluntary decisions.

Lastly, our results extend CPP findings from perceptual tasks to voluntary, internally driven choices, including simple value-based decisions. Although this lends support to the view that the CPP acts as a domain-general correlate of evidence accumulation, previous reports of CPP-like signals in value-based (Pisauro et al., 2017) and memory-based decisions (van Ede & Nobre, 2024; van Vugt et al., 2019) have warranted careful interpretation due to methodological limitations in these studies (O’Connell et al., 2025). To validate prior CPP reports while addressing methodological concerns raised, the present analysis framework could be applied to value- and memory-based decision datasets, as well as other domains (e.g., food preference and moral decision-making) to more decisively adjudicate whether the CPP consistently tracks evidence accumulation beyond perceptual and voluntary domains.

### 4.5 Conclusion

This study extends our current understanding of how voluntary decisions evolve and are converted into action. Despite prior reports of anatomical and qualitative differences between voluntary and forced actions, we provide neural evidence that voluntary decisions, like forced decisions, unfold as a noisy evidence accumulation process that terminates at a stereotypical boundary. Crucially, the presence of canonical CPP dynamics in voluntary and simple value-based decisions suggests that the CPP is not exclusive to perceptual decision making. The LHRP did not show clear evidence of an evolving decision signal under matched motor demands, and may instead reflect late effector-specific preparation.

Together, these findings point to common decision commitment mechanisms and motor-gating mechanisms between internally and externally driven decisions.

## Data and Code Availability Statement

All code and data used for the experiment and analyses described in this manuscript are available at http://osf.io/t7z4h.

## CRediT authorship contribution statement

**Lauren C. Fong**: Writing – review & editing, Writing – original draft, Visualisation, Methodology, Formal analysis, Data curation, Conceptualisation. **Paul M. Garrett**: Writing – review, & editing, Formal analysis. **Philip L. Smith**: Writing – review & editing, Supervision. **Robert Hester**: Writing – review & editing, Supervision. **Stefan Bode**: Writing – review & editing, Supervision, Funding acquisition, Conceptualisation. **Daniel Feuerriegel**: Writing – review & editing, Visualisation, Supervision, Resources, Project administration, Methodology, Funding acquisition, Formal analysis, Conceptualisation.

## Ethical statement

The experiment was approved by the Human Research Ethics Committee (ID 27017) of the University of Melbourne.

## Acknowledgements

We thank our laboratory members, Jie Sun in particular, and Redmond O’Connell, Simon Kelly, Elaine Corbett, Elisabeth Parés-Pujolràs, and Silvia Seghezzi for helpful discussions relating to this paper.

