## Supplementary Materials for "Tracing the neural trajectories of evidence accumulation and motor preparation processes during voluntary decisions"

Saadiyat Marina District - Abu Dhabi - United Arab Emirates

### Change-of-Mind Decision RTs

On average, voluntary decisions were slower than forced decisions in both the initial (as reported in the main results) and subsequent change-of-mind decision phases. Voluntary "stayed" decisions ($M$ = 1.28 s) were 0.21 s slower than forced "stayed" decisions ($M$ = 1.07 s), $t(48) = 4.18$, $p < .001$, $d_z = 0.60$, 95% CI [0.11, 0.32]. Voluntary "switched" decisions ($M$ = 1.34 s) were 0.17 s slower than forced "switched" decisions ($M$ = 1.17 s), $t(48) = 3.79$, $p < .001$, $d_z = 0.54$, 95% CI [0.08, 0.26].

Within voluntary subsequent decisions, "switched" and "stayed" decisions did not differ reliably in RTs, $t(48) = -1.90$, $p = .064$, $d_z = -0.27$, mean difference = –0.06 s, 95% CI [–0.11, 0.003]. However, within forced subsequent decisions, "switched" decisions were –0.10 s slower than "stayed" decisions, $t(48) = -3.48$, $p = .001$, $d_z = -0.50$, 95% CI [–0.16, –0.04]. This suggests a switch cost associated with decision updating but only for forced decisions.

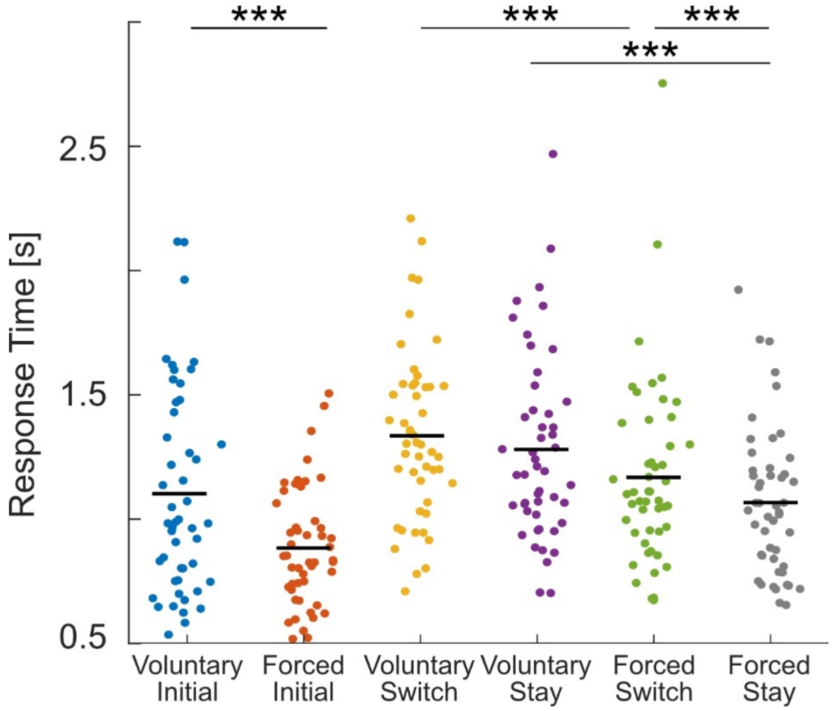


***Supplementary Figure S1.*** Participant (dots) and group (horizontal black lines) mean response times by decision type. Lines with asterisks denote statistically significant response time differences (*** denotes p < .001).

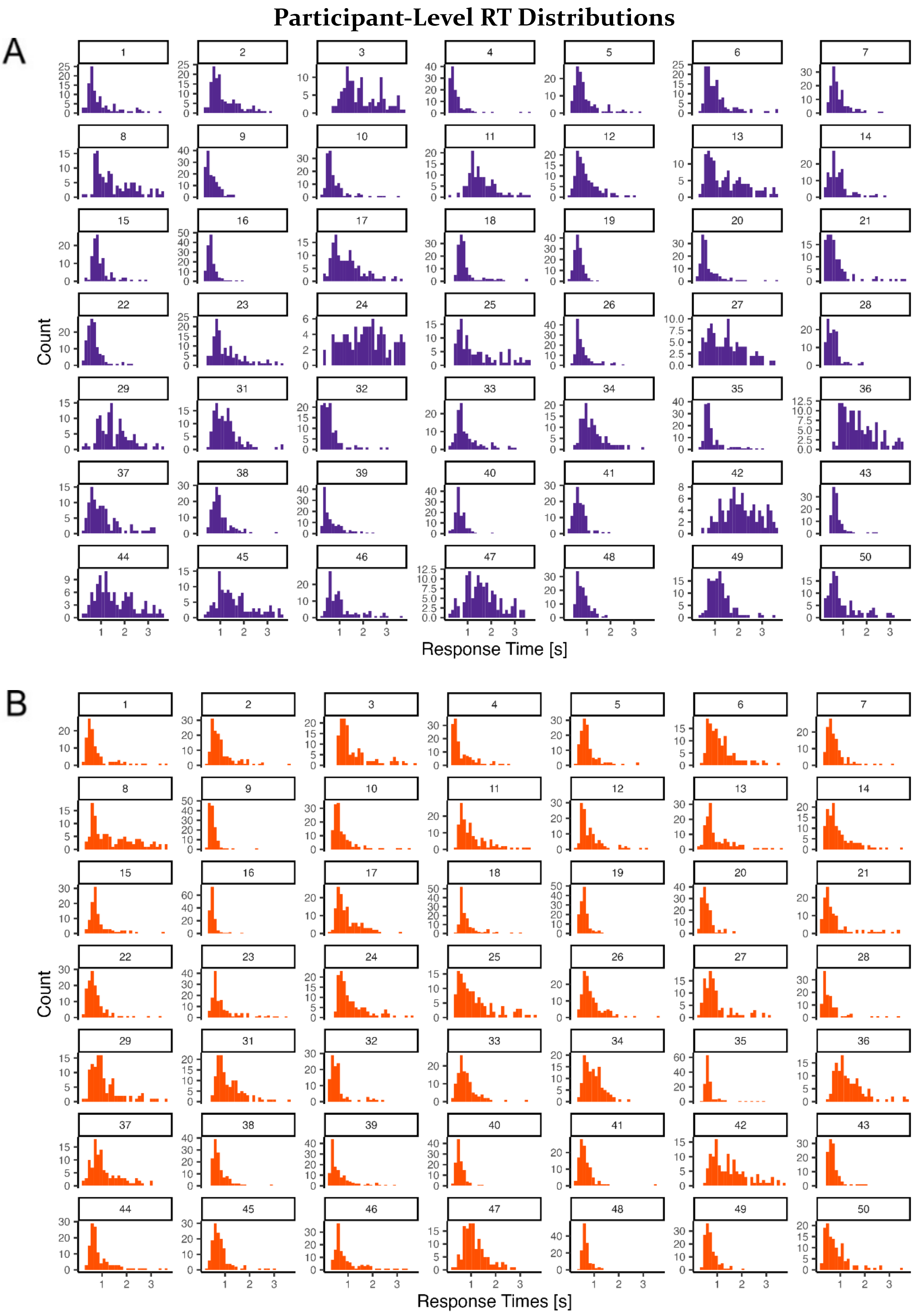


***Supplementary Figure S2.*** Participant-level response time distributions for A) voluntary and B) forced decisions. Each panel shows RT distributions for each participant that are positively skewed across most participants.

### Choice Behaviour

For each participant's voluntary initial decision trials, we computed the following and summarised them in Supplementary Table 1:

- *N* Chosen: the number of trials in which each colour was chosen
- *N* Presented: the number of trials in which each colour was presented as an option
- Mean RT: the mean RT on voluntary trials in which each colour was chosen.
- Frequency Ranking: rank ordering of colours based on *N* Chosen (most frequent = 1; least frequent = 4).
- RT Ranking: rank ordering of colours by Mean RT for each colour (fastest = 1; NA = never chosen)
- Self-Reported Ranking: participants' self-reported colour preference ranking collected at the end of the experiment
- Self-Reported Strategy: participants' self-reported choice strategies collected at the end of the experiment

Supplementary Table 1 shows several patterns consistent with deliberate, goal- or preference-guided behaviour, some of which include: (1) close alignment between self-reported preference rankings and choice-frequency rankings (e.g., IDs 2, 10, 31, 32), consistent with preference- or goal-driven deliberation, (2) fastest RTs for the highest-ranked colours (e.g., IDs 23, 31, 44, 45) or for the lowest-ranked colours (e.g., IDs 9, 15, 22, 46), (4) slowest RTs for the lowest-ranked colours (e.g., IDs 12, 27, 48), consistent with increased deliberation when choosing a less-preferred option, (5) relatively similar choice frequencies across colours among participants who reported a diversity-maximisation strategy (e.g., IDs 33, 44, 47), (6) relatively similar RT profiles when choice frequencies were also relatively balanced across colours (e.g., IDs 16, 28, 29, 40), (7) strong avoidance of lowest-ranked colours and very high choice frequency for highest-ranked colours (e.g., IDs 19, 26, 35), suggesting stable decision strategies. We note that these examples are not an exhaustive list of all strategies but rather illustrate that many participants showed stable patterns consistent with goal- or preference-guided choice, while others showed more heterogeneous patterns—an expected feature of voluntary decision-making where choice can be guided by endogenous factors that may vary within and across individuals, some of which may also not be easily available for conscious reflection.

**Supplementary Table 1**

*Summary of Participant-Level Colour Choice Behaviour and Self-Reported Preferences and Choice Strategies.*

| ID | Chosen Colour | *N* Chosen | *N* Presented | Mean RT | Frequency Ranking | Self-Reported Ranking | RT Ranking | Self-Reported Strategy |
|---|---|---|---|---|---|---|---|---|
| 1 | Blue | 38 | 70 | 1.04 | 1 | 1 | 3 | Random |
| | Green | 37 | 71 | 1.02 | 2 | 4 | 2 | |
| | Orange | 33 | 70 | 0.83 | 3 | 3 | 1 | |
| | Pink | 32 | 69 | 1.08 | 4 | 2 | 4 | |
| 2 | Blue | 45 | 71 | 1.14 | 1 | 1 | 2 | Momentary |
| | Green | 30 | 71 | 1.15 | 3 | 3 | 3 | |
| | Orange | 28 | 72 | 1.05 | 4 | 4 | 1 | |
| | Pink | 40 | 72 | 1.24 | 2 | 2 | 4 | |
| 3 | Blue | 30 | 62 | 1.96 | 2 | 3 | 3 | Momentary |
| | Green | 39 | 63 | 2.09 | 1 | 1 | 4 | |
| | Orange | 28 | 61 | 1.86 | 3 | 2 | 2 | |
| | Pink | 26 | 60 | 1.85 | 4 | 4 | 1 | |
| 4 | Blue | 45 | 67 | 0.53 | 2 | 2 | 3 | Preference |
| | Green | 62 | 66 | 0.60 | 1 | 1 | 4 | |
| | Orange | 3 | 68 | 0.47 | 4 | 4 | 2 | |
| | Pink | 23 | 65 | 0.44 | 3 | 3 | 1 | |
| 5 | Blue | 41 | 68 | 0.98 | 2 | 2 | 4 | Momentary |
| | Green | 27 | 68 | 0.83 | 3 | 4 | 1 | |
| | Orange | 25 | 70 | 0.91 | 4 | 3 | 2 | |
| | Pink | 44 | 68 | 0.91 | 1 | 1 | 3 | |
| 6 | Blue | 46 | 72 | 0.97 | 1 | 1 | 1 | Diverse Range |
| | Green | 37 | 72 | 1.24 | 3 | 3 | 4 | |
| | Orange | 19 | 71 | 1.03 | 4 | 4 | 2 | |
| | Pink | 41 | 71 | 1.07 | 2 | 2 | 3 | |
| 7 | Blue | 57 | 72 | 0.90 | 1 | 1 | 3 | Location |
| | Green | 26 | 72 | 0.78 | 4 | 4 | 1 | |
| | Orange | 30 | 72 | 0.88 | 3 | 2 | 2 | |
| | Pink | 31 | 72 | 0.92 | 2 | 3 | 4 | |
| 8 | Blue | 38 | 58 | 1.44 | 2 | 1 | 1 | Momentary |
| | Green | 41 | 61 | 1.61 | 1 | 2 | 3 | |
| | Orange | 26 | 64 | 1.71 | 3 | 4 | 4 | |
| | Pink | 18 | 63 | 1.55 | 4 | 3 | 2 | |
| 9 | Blue | 37 | 71 | 0.70 | 2 | 3 | 4 | Preference |
| | Green | 34 | 69 | 0.69 | 3 | 2 | 3 | |
| | Orange | 56 | 72 | 0.62 | 1 | 1 | 2 | |
| | Pink | 13 | 68 | 0.49 | 4 | 4 | 1 | |

| | | | | | | | | |
|---|---|---|---|---|---|---|---|---|
| 10 | Blue | 44 | 71 | 0.73 | 2 | 2 | 2 | Momentary |
| | Green | 12 | 71 | 0.63 | 4 | 4 | 1 | |
| | Orange | 17 | 72 | 0.79 | 3 | 3 | 3 | |
| | Pink | 70 | 72 | 0.92 | 1 | 1 | 4 | |
| 11 | Blue | 41 | 67 | 1.76 | 1 | 3 | 4 | Avoid Sequential Choices |
| | Green | 25 | 64 | 1.60 | 4 | 4 | 3 | |
| | Orange | 31 | 58 | 1.40 | 2 | 2 | 1 | |
| | Pink | 29 | 63 | 1.57 | 3 | 1 | 2 | |
| 12 | Blue | 44 | 57 | 1.02 | 1 | 1 | 2 | Momentary |
| | Green | 18 | 59 | 1.33 | 4 | 4 | 4 | |
| | Orange | 23 | 55 | 0.71 | 3 | 3 | 1 | |
| | Pink | 30 | 59 | 1.05 | 2 | 2 | 3 | |
| 13 | Blue | 22 | 65 | 1.59 | 4 | 3 | 3 | Random |
| | Green | 39 | 67 | 1.70 | 1 | 1 | 4 | |
| | Orange | 37 | 70 | 1.29 | 3 | 4 | 1 | |
| | Pink | 37 | 68 | 1.35 | 2 | 2 | 2 | |
| 14 | Blue | 45 | 69 | 0.87 | 2 | 2 | 4 | Preference |
| | Green | 29 | 69 | 0.80 | 3 | 3 | 2 | |
| | Orange | 12 | 65 | 0.73 | 4 | 4 | 1 | |
| | Pink | 48 | 65 | 0.81 | 1 | 1 | 3 | |
| 15 | Blue | 40 | 69 | 1.05 | 2 | 2 | 4 | Preference |
| | Green | 3 | 66 | 0.80 | 4 | 4 | 1 | |
| | Orange | 24 | 64 | 1.03 | 3 | 3 | 3 | |
| | Pink | 66 | 67 | 0.91 | 1 | 1 | 2 | |
| 16 | Blue | 36 | 72 | 0.62 | 2 | 2 | 1 | Preference |
| | Green | 40 | 72 | 0.67 | 1 | 1 | 4 | |
| | Orange | 35 | 72 | 0.67 | 3 | 4 | 3 | |
| | Pink | 33 | 72 | 0.62 | 4 | 3 | 2 | |
| 17 | Blue | 42 | 68 | 1.36 | 1 | 2 | 3 | Random |
| | Green | 26 | 68 | 1.29 | 4 | 4 | 2 | |
| | Orange | 33 | 68 | 1.24 | 3 | 3 | 1 | |
| | Pink | 35 | 68 | 1.40 | 2 | 1 | 4 | |
| 18 | Blue | 51 | 72 | 0.99 | 2 | 2 | 3 | Momentary |
| | Green | 21 | 71 | 1.10 | 3 | 3 | 4 | |
| | Orange | 15 | 71 | 0.79 | 4 | 4 | 1 | |
| | Pink | 56 | 72 | 0.92 | 1 | 1 | 2 | |
| 19 | Blue | 51 | 72 | 0.66 | 2 | 2 | 3 | Diverse Range |
| | Green | 30 | 72 | 0.66 | 3 | 3 | 2 | |
| | Orange | 0 | 72 | NA | 4 | 4 | NA | |
| | Pink | 63 | 72 | 0.64 | 1 | 1 | 1 | |
| 20 | Blue | 35 | 71 | 0.91 | 3 | 2 | 4 | Preference |
| | Green | 36 | 71 | 0.69 | 2 | 4 | 2 | |

| | | | | | | | | |
|---|---|---|---|---|---|---|---|---|
| | Orange | 33 | 71 | 0.56 | 4 | 3 | 1 | |
| | Pink | 38 | 71 | 0.87 | 1 | 1 | 3 | |
| 21 | Blue | 41 | 62 | 0.90 | 1 | 1 | 4 | Random |
| | Green | 21 | 58 | 0.69 | 3 | 3 | 2 | |
| | Orange | 22 | 61 | 0.66 | 4 | 4 | 1 | |
| | Pink | 36 | 59 | 0.80 | 2 | 2 | 3 | |
| 22 | Blue | 66 | 72 | 0.73 | 1 | 1 | 4 | Preference |
| | Green | 13 | 71 | 0.59 | 4 | 4 | 1 | |
| | Orange | 17 | 72 | 0.67 | 3 | 3 | 2 | |
| | Pink | 47 | 71 | 0.73 | 2 | 2 | 3 | |
| 23 | Blue | 63 | 70 | 1.14 | 1 | 1 | 1 | Preference |
| | Green | 32 | 70 | 1.72 | 3 | 3 | 4 | |
| | Orange | 34 | 71 | 1.19 | 2 | 2 | 2 | |
| | Pink | 12 | 71 | 1.63 | 4 | 4 | 3 | |
| 24 | Blue | 22 | 48 | 2.45 | 4 | 3 | 4 | Preference |
| | Green | 25 | 53 | 1.98 | 3 | 4 | 1 | |
| | Orange | 26 | 53 | 2.00 | 2 | 1 | 2 | |
| | Pink | 23 | 38 | 2.02 | 1 | 2 | 3 | |
| 25 | Blue | 30 | 61 | 1.34 | 3 | 3 | 1 | Random |
| | Green | 28 | 59 | 1.44 | 4 | 2 | 3 | |
| | Orange | 33 | 65 | 1.65 | 2 | 1 | 4 | |
| | Pink | 31 | 59 | 1.36 | 1 | 4 | 2 | |
| 26 | Blue | 25 | 72 | 0.77 | 3 | 3 | 1 | Preference |
| | Green | 47 | 72 | 0.86 | 2 | 2 | 2 | |
| | Orange | 72 | 72 | 0.87 | 1 | 1 | 3 | |
| | Pink | 0 | 72 | NA | 4 | 4 | NA | |
| 27 | Blue | 38 | 64 | 1.48 | 2 | 2 | 3 | Momentary |
| | Green | 35 | 62 | 1.42 | 3 | 3 | 2 | |
| | Orange | 9 | 63 | 1.91 | 4 | 4 | 4 | |
| | Pink | 43 | 61 | 1.40 | 1 | 1 | 1 | |
| 28 | Blue | 26 | 60 | 0.57 | 3 | 1 | 3 | Preference |
| | Green | 24 | 62 | 0.56 | 4 | 3 | 2 | |
| | Orange | 33 | 60 | 0.68 | 2 | 2 | 4 | |
| | Pink | 36 | 56 | 0.55 | 1 | 4 | 1 | |
| 29 | Blue | 35 | 61 | 1.56 | 2 | 1 | 2 | Preference |
| | Green | 37 | 62 | 1.57 | 1 | 2 | 3 | |
| | Orange | 26 | 67 | 1.67 | 4 | 3 | 4 | |
| | Pink | 30 | 66 | 1.53 | 3 | 4 | 1 | |
| 31 | Blue | 44 | 72 | 1.21 | 2 | 2 | 2 | Preference |
| | Green | 11 | 70 | 1.47 | 4 | 4 | 3 | |
| | Orange | 19 | 71 | 1.51 | 3 | 3 | 4 | |
| | Pink | 68 | 71 | 1.15 | 1 | 1 | 1 | |

| | | | | | | | | |
|---|---|---|---|---|---|---|---|---|
| 32 | Blue | 40 | 63 | 0.62 | 2 | 2 | 3 | Preference |
| | Green | 1 | 66 | 0.52 | 4 | 4 | 1 | |
| | Orange | 27 | 63 | 0.54 | 3 | 3 | 2 | |
| | Pink | 60 | 64 | 0.65 | 1 | 1 | 4 | |
| 33 | Blue | 37 | 64 | 0.92 | 2 | 1 | 2 | Diverse Range |
| | Green | 22 | 65 | 1.20 | 4 | 3 | 4 | |
| | Orange | 43 | 66 | 0.90 | 1 | 2 | 1 | |
| | Pink | 27 | 63 | 0.96 | 3 | 4 | 3 | |
| 34 | Blue | 37 | 68 | 1.27 | 2 | 1 | 3 | Avoid Sequential Choices |
| | Green | 29 | 70 | 1.19 | 4 | 3 | 1 | |
| | Orange | 32 | 68 | 1.24 | 3 | 2 | 2 | |
| | Pink | 40 | 70 | 1.29 | 1 | 4 | 4 | |
| 35 | Blue | 77 | 77 | 0.83 | 1 | 1 | 1 | Preference |
| | Green | 25 | 71 | 0.99 | 3 | 3 | 3 | |
| | Orange | 46 | 74 | 0.96 | 2 | 2 | 2 | |
| | Pink | 0 | 74 | NA | 4 | 4 | NA | |
| 36 | Blue | 33 | 66 | 1.75 | 3 | 2 | 4 | Preference |
| | Green | 37 | 68 | 1.67 | 2 | 1 | 3 | |
| | Orange | 25 | 66 | 1.41 | 4 | 4 | 1 | |
| | Pink | 37 | 64 | 1.65 | 1 | 3 | 2 | |
| 37 | Blue | 42 | 62 | 1.20 | 1 | 1 | 3 | Preference |
| | Green | 24 | 64 | 0.92 | 3 | 3 | 1 | |
| | Orange | 21 | 63 | 1.02 | 4 | 4 | 2 | |
| | Pink | 40 | 65 | 1.24 | 2 | 2 | 4 | |
| 38 | Blue | 42 | 72 | 0.91 | 2 | 2 | 2 | Preference |
| | Green | 29 | 72 | 1.03 | 3 | 3 | 3 | |
| | Orange | 1 | 72 | 0.67 | 4 | 4 | 1 | |
| | Pink | 72 | 72 | 1.06 | 1 | 1 | 4 | |
| 39 | Blue | 21 | 70 | 0.90 | 3 | 3 | 4 | Preference |
| | Green | 58 | 72 | 0.62 | 1 | 1 | 2 | |
| | Orange | 46 | 67 | 0.67 | 2 | 2 | 3 | |
| | Pink | 14 | 69 | 0.48 | 4 | 4 | 1 | |
| 40 | Blue | 40 | 72 | 0.69 | 1 | 1 | 2 | Preference |
| | Green | 37 | 72 | 0.66 | 2 | 2 | 1 | |
| | Orange | 34 | 72 | 0.71 | 3 | 3 | 4 | |
| | Pink | 33 | 72 | 0.70 | 4 | 4 | 3 | |
| 41 | Blue | 39 | 65 | 0.66 | 1 | 2 | 1 | Salience |
| | Green | 27 | 65 | 0.68 | 4 | 4 | 2 | |
| | Orange | 35 | 65 | 0.75 | 2 | 3 | 4 | |
| | Pink | 29 | 65 | 0.73 | 3 | 1 | 3 | |
| 42 | Blue | 32 | 54 | 2.10 | 1 | 2 | 1 | Momentary |
| | Green | 18 | 44 | 2.13 | 3 | 3 | 3 | |

| | | | | | | | | |
|---|---|---|---|---|---|---|---|---|
| | Orange | 18 | 47 | 2.14 | 4 | 4 | 4 | |
| | Pink | 29 | 49 | 2.11 | 2 | 1 | 2 | |
| 43 | Blue | 39 | 70 | 0.73 | 1 | 2 | 2 | Preference |
| | Green | 36 | 72 | 0.82 | 3 | 4 | 3 | |
| | Orange | 40 | 72 | 0.73 | 2 | 1 | 1 | |
| | Pink | 27 | 70 | 0.86 | 4 | 3 | 4 | |
| 44 | Blue | 34 | 70 | 1.58 | 3 | 3 | 2 | Diverse Range |
| | Green | 43 | 67 | 1.50 | 1 | 1 | 1 | |
| | Orange | 25 | 66 | 1.72 | 4 | 2 | 4 | |
| | Pink | 32 | 65 | 1.62 | 2 | 4 | 3 | |
| 45 | Blue | 39 | 61 | 1.59 | 2 | 2 | 2 | Salience |
| | Green | 42 | 65 | 1.56 | 1 | 1 | 1 | |
| | Orange | 16 | 62 | 1.98 | 4 | 4 | 4 | |
| | Pink | 25 | 56 | 1.69 | 3 | 3 | 3 | |
| 46 | Blue | 44 | 65 | 1.06 | 2 | 2 | 3 | Preference |
| | Green | 35 | 67 | 1.13 | 3 | 3 | 4 | |
| | Orange | 5 | 65 | 0.69 | 4 | 4 | 1 | |
| | Pink | 46 | 63 | 1.06 | 1 | 1 | 2 | |
| 47 | Blue | 30 | 67 | 1.69 | 4 | 3 | 3 | Diverse Range |
| | Green | 37 | 69 | 1.60 | 1 | 1 | 2 | |
| | Orange | 34 | 69 | 1.47 | 3 | 4 | 1 | |
| | Pink | 36 | 69 | 1.77 | 2 | 2 | 4 | |
| 48 | Blue | 48 | 72 | 0.82 | 1 | 2 | 2 | Preference |
| | Green | 22 | 69 | 0.95 | 4 | 4 | 4 | |
| | Orange | 23 | 69 | 0.86 | 3 | 3 | 3 | |
| | Pink | 48 | 72 | 0.81 | 2 | 1 | 1 | |
| 49 | Blue | 49 | 72 | 1.16 | 1 | 1 | 1 | Preference |
| | Green | 25 | 72 | 1.19 | 3 | 4 | 2 | |
| | Orange | 22 | 72 | 1.32 | 4 | 3 | 4 | |
| | Pink | 48 | 72 | 1.20 | 2 | 2 | 3 | |
| 50 | Blue | 27 | 64 | 1.10 | 4 | 4 | 4 | Random |
| | Green | 29 | 67 | 0.87 | 3 | 1 | 1 | |
| | Orange | 34 | 64 | 0.96 | 2 | 2 | 2 | |
| | Pink | 41 | 67 | 0.97 | 1 | 3 | 3 | |

### Mixed Effects Regression Model Structures and Coefficients for Analyses of Pre-Response Amplitudes, Slopes, and Response Times

As described in the EEG Analysis section of the Methods, we fitted linear mixed-effects models (LMMs) using the lme4 package (Bates et al., 2015) to relate response times (RTs) to pre-response amplitude and slope measures for each signal – the centro-parietal positivity (CPP), Mu/Beta (MB) and left hemisphere readiness potential (LHRP). By default, we used a model with participant intercepts and random slopes for RT, allowing the strength of the RT–EEG measure relationship to vary across participants. If the model produced convergence or singular fit issues, we removed the random slopes for RT but retained the random intercepts.

We compared models with and without the fixed effect of RT using maximum likelihood and Chi-Square likelihood-ratio tests within matched random-effects structures as follows and report the results in Supplementary Table 2 below:

- m4: $EEG_{measure} \sim 1 + RT + (1 + RT \mid pID)$ vs m3: $EEG_{measure} \sim 1 + (1 + RT \mid pID)$
- m2: $EEG_{measure} \sim 1 + RT + (1 \mid pID)$ vs m1: $EEG_{measure} \sim 1 + (1 \mid pID)$

These likelihood ratio tests serve as a transparency check to test whether adding a fixed effect of RT meaningfully improves model fit, holding the random-effects structure constant. A significant LRT result indicates that including RT as a fixed effect improves fit, supporting an RT–EEG measure relationship that is reliably non-zero at the group level, given the specified random-effects structure. By contrast, a non-significant LRT result suggests that the data do not strongly support a consistent group-level RT effect for that signal, even if individuals may still vary (as captured by random effects).

Because LRTs and coefficient tests answer slightly different questions (model-fit improvement vs. estimating a parameter within a given model), occasional differences can occur for small or noisy effects. We therefore use the LRT outcomes mainly to qualify how strongly the dataset supports adding RT as an explanatory predictor, while the reported fixed-effect coefficients provide the estimated direction and size of the RT–EEG measure associations.

**Supplementary Table 2**

*Chi-Square Likelihood Ratio Tests for Group-Level Model Comparisons.*

| Signal | Decision Type | ERP Measure | Chi-square | df | p | Comparison Test | Best Model |
|---|---|---|---|---|---|---|---|
| CPP | Voluntary | Slope | 20.52 | 1 | <.001*** | m1 vs m2 | m4 |
| | | | 13.74 | 1 | <.001*** | m3 vs m4 | |
| | | Amplitude | 4.66 | 1 | .031* | m1 vs m2 | m3 |
| | | | 1.30 | 1 | .254 | m3 vs m4 | |
| | Forced | Slope | 18.06 | 1 | <.001*** | m1 vs m2 | m4 |
| | | | 10.11 | 1 | .002** | m3 vs m4 | |
| | | Amplitude | 2.56 | 1 | .109 | m1 vs m2 | m1 |
| | | | 1.44 | 1 | .230 | m3 vs m4 | |
| MB | Voluntary | Slope | 2.16 | 1 | .141 | m1 vs m2 | m1 |
| | | | 1.98 | 1 | .160 | m3 vs m4 | |
| | | Amplitude | 0.20 | 1 | .653 | m1 vs m2 | m3 |
| | | | 0.19 | 1 | .664 | m3 vs m4 | |
| | Forced | Slope | 0.01 | 1 | .909 | m1 vs m2 | m1 |
| | | | 0.02 | 1 | .893 | m3 vs m4 | |
| | | Amplitude | 0.51 | 1 | .475 | m1 vs m2 | m3 |
| | | | 0.56 | 1 | .452 | m3 vs m4 | |
| LHRP | Voluntary | Slope | 7.23 | 1 | .007** | m1 vs m2 | m4 |
| | | | 5.33 | 1 | .021* | m3 vs m4 | |
| | | Amplitude | 0.32 | 1 | .571 | m1 vs m2 | m1 |
| | | | 0.10 | 1 | .754 | m3 vs m4 | |
| | Forced | Slope | 1.55 | 1 | .213 | m1 vs m2 | m3 |
| | | | 1.06 | 1 | .303 | m3 vs m4 | |
| | | Amplitude | 14.35 | 1 | <.001*** | m1 vs m2 | m4 |
| | | | 6.44 | 1 | .011* | m3 vs m4 | |

*Note*. Models denoted in red produced a singular fit and failed to converge.
*p <.05 **p <.01 ***p <.001

**Supplementary Table 3**

Linear Mixed-Effects Model Coefficients for Predicting CPP Pre-Response Amplitudes and Slopes from Reaction Times

| Decision Type | CPP measure | Model | Estimate | SE | t | p |
|---|---|---|---|---|---|---|
| Voluntary | Amplitude | m2 | 1.78 | 0.82 | 2.16 | .031* |
| | | **m4** | **1.55** | **1.34** | **1.15** | **.255** |
| | Slope | m2 | -0.02 | <.01 | -4.53 | <.001*** |
| | | **m4** | **-0.02** | **<.01** | **-4.03** | **<.001*** ** |
| Forced | Amplitude | **m2** | **1.20** | **0.75** | **1.60** | **.109** |
| | | m4 | 0.72 | 1.07 | 0.68 | .502 |
| | Slope | m2 | -0.02 | 0.005 | -4.25 | <.001*** |
| | | **m4** | **-0.02** | **.01** | **-3.34** | **.002** ** |

*Note.* m2 denotes the random intercept model. m4 denotes the random slopes model. Rows in red indicate results from models that failed to converge or produced a singular fit. Rows in bold indicate results reported in the main manuscript. *p <.05 **p <.01 ***p <.001

**Supplementary Table 4**

Linear Mixed-Effects Model Coefficients for Predicting MB Pre-Response Amplitudes and Slopes from Reaction Times

| Decision Type | MB measure | Model | Estimate | SE | t | p |
|---|---|---|---|---|---|---|
| Voluntary | Amplitude | m2 | 0.03 | 0.07 | 0.45 | .653 |
| | | **m4** | **0.03** | **0.07** | **0.43** | **.665** |
| | Slope | **m2** | **0.002** | **0.001** | **1.47** | **<.001*** ** |
| | | m4 | 0.002 | 0.001 | 1.35 | <.001*** |
| Forced | Amplitude | m2 | -0.05 | 0.07 | -0.71 | .475 |
| | | **m4** | **-0.06** | **0.09** | **-0.75** | **.456** |
| | Slope | **m2** | **0.0001** | **0.001** | **0.11** | **.141** |
| | | m4 | 0.0002 | 0.001 | 0.14 | .183 |

*Note.* m2 denotes the random intercept model. m4 denotes the random slopes model. Rows in red indicate results from models that failed to converge or produced a singular fit. Rows in bold indicate results reported in the main manuscript. *p <.05 **p <.01 ***p <.001

**Supplementary Table 5**

Linear Mixed-Effects Model Coefficients for Predicting LHRP Pre-Response Amplitudes and Slopes from Reaction Times

| Decision Type | LHRP measure | Model | Estimate | SE | t | p |
|---|---|---|---|---|---|---|
| Voluntary | Amplitude | **m2** | **-0.56** | **0.99** | **-0.57** | **.571** |
| | | m4 | -0.36 | 1.16 | -0.31 | .754 |
| | Slope | m2 | 0.01 | <.01 | 2.69 | .007** |
| | | **m4** | **0.01** | **<.01** | **2.42** | **.019*** |
| Forced | Amplitude | m2 | -3.44 | 0.91 | -3.79 | <.001*** |
| | | **m4** | **-3.53** | **1.34** | **-2.62** | **.012*** |
| | Slope | m2 | -0.01 | <.01 | -1.25 | .213 |
| | | **m4** | **-0.01** | **0.01** | **-1.03** | **.306** |

*Note.* m2 denotes the random intercept model. m4 denotes the random slopes model. Rows in red indicate results from models that failed to converge or produced a singular fit. Rows in bold indicate results reported in the main manuscript. *p <.05 **p <.01 ***p <.001

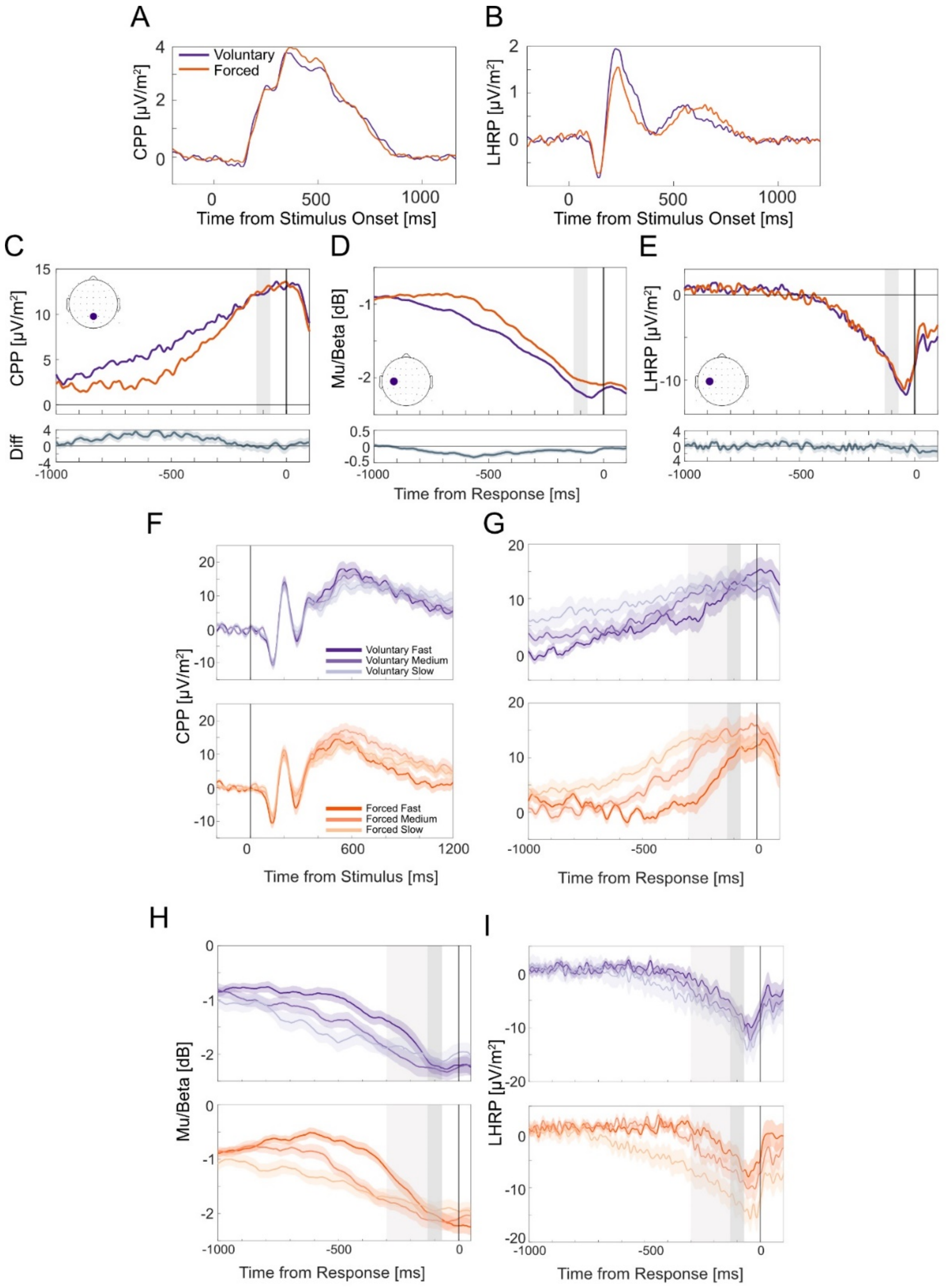


***Supplementary Figure S3.*** CPP, Mu/Beta, and LHRP amplitude waveforms. A – B) RIDE-derived S (stimulus-locked) subcomponent group-average waveforms at electrode Pz (A) and C3 (B), reflecting ERP waveform contributions that are strictly locked to stimulus onset. C) Group-averaged CPP, D) MB amplitudes, and E) LHRP for voluntary (purple) and forced (orange) waveforms with accompanying difference waves (and shaded standard error regions) at each time point. Plotted waveforms are for data where the S component was not subtracted and the overlap of stimulus-locked waveforms was not corrected for. Grey areas represent the pre-response windows between -130 to -70 ms. F) Group averaged CPP relative to the stimulus onset for voluntary (purple) and forced (orange) decisions split by fast, medium, and slow RT bins, with shading denoting standard errors. G) Group averaged CPP, H) MB amplitudes, and I) LHRP for voluntary (purple) and forced (orange) decisions relative to the response split by fast, medium, and slow RT bins, with shading denoting standard errors. Dark grey areas denote the -130 to -70 ms pre-response amplitude window. Light grey areas denote the -300 to -70 ms time slope window. Overall, the response-locked waveforms in Figures G-I look similar to the data where the S component had been subtracted.

### Supplementary Material References

Bates, D., Maechler, M., Bolker, B., & Walker, S. (2015). Fitting linear mixed-effects models using lme4. *Journal of Statistical Software, 67*(1), 1-48.
